# Towards Reliable Diabetes Prediction: Innovations in Data Engineering and Machine Learning Applications

**DOI:** 10.1101/2024.07.14.603436

**Authors:** Md. Alamin Talukder, Md. Manowarul Islam, Md Ashraf Uddin, Mohsin Kazi, Majdi Khalid, Arnisha Akhter, Mohammad Ali Moni

## Abstract

**Objective:** Diabetes is a metabolic disorder that causes the risk of stroke, heart disease, kidney failure, and other long-term complications because diabetes generates excess sugar in the blood. Machine learning (ML) models can aid in diagnosing diabetes at the primary stage. So, we need an efficient machine learning model to diagnose diabetes accurately.

**Methods:** In this paper, an effective data preprocessing pipeline has been implemented to process the data and random oversampling to balance the data, handling the imbalance distributions of the observational data more sophisticatedly. We used four different diabetes datasets to conduct our experiments. Several ML algorithms were used to determine the best models to predict diabetes faultlessly.

**Results:** The performance analysis demonstrates that among all ML algorithms, RF surpasses the current works with an accuracy rate of 86% and 98.48% for dataset-1 and dataset-2; XGB and DT surpass with an accuracy rate of 99.27% and 100% for dataset-3 and dataset-4 respectively. Our proposal can increase accuracy by 12.15% compared to the model without preprocessing.

**Conclusions:** This excellent research finding indicates that the proposed models might be employed to produce more accurate diabetes predictions to supplement current preventative interventions to reduce the incidence of diabetes and its associated costs.

## Introduction

Diabetes mellitus (DM) is a chronic disorder that affects carbohydrate, protein, and fat metabolism, leading to abnormal blood glucose levels (*1*). It is classified into two main types: type 1 and type 2 diabetes (*2*). Type 1 diabetes typically occurs in children but can manifest in adults, particularly in their late thirties and early forties. Patients with type 1 diabetes are usually not obese and often present with a life-threatening condition known as diabetic ketoacidosis (*3*). The etiology of type 1 diabetes involves damage to pancreatic cells due to environmental or infectious agents, triggering an autoimmune response against *β*-cells. Autoimmunity is considered the primary factor in the pathophysiology of type 1 diabetes. Type 1 diabetes is also associated with other autoimmune diseases (*4*). On the other hand, type 2 diabetes has a distinct pathophysiology and etiology, characterized by a combination of low insulin production and insulin resistance. Obesity, physical inactivity, poor diet, and urbanization contribute to the rising prevalence of type 2 diabetes (*5*). Dysfunction of *β*-cells plays a crucial role in progressing from prediabetes to diabetes. Despite their differing pathophysiology, both types of diabetes share similar complications, including macrovascular and microvascular complications (*6*). It is a severe chronic disease (*7*), associated with various consequences and increased mortality (*8*).

Health regulations emphasize regular screenings for individuals with diabetes risk factors (*9*), highlighting the importance of timely identification and intervention. Preventive measures are crucial alongside diabetes care (*10*). Early diagnosis and lifestyle modifications, such as healthy eating and exercise, can reduce the progression from impaired glucose tolerance to prediabetes (*11*). Technology, particularly machine learning (ML), has gained popularity for early detection and prevention in healthcare (*12–15*). ML in diabetes management offers a promising avenue for predictive modeling. By analyzing vast datasets encompassing patient demographics, medical history, and lifestyle factors, ML algorithms can predict the likelihood of diabetes onset or progression with remarkable accuracy. These models not only assist in early detection but also empower healthcare providers to tailor personalized interventions, ultimately mitigating complications and improving patient outcomes (*16, 17*)

Several ML algorithms have been introduced for diabetes detection, offering benefits such as low computation costs, robustness, and high performance (*18*). For instance, researchers have utilized classifiers like Nave Bayes (NB), Decision Tree (DT), Adaptive Boosting (Adaboost) and Random Forest (RF) for diabetes prediction (*19*), while models such as Generalized Linear Models with Elastic Net Regularization (Glmnet), RF, Extreme Gradient Boosting (XGBoost), and Light Gradient Boosting Machine (LightGBM) have been explored for predicting Type-2 diabetes (*20*). According to recent projections, the prevalence of diabetes is expected to rise significantly, imposing a substantial burden on healthcare systems worldwide (*21*). Early detection and effective management of diabetes are crucial for preventing complications and improving patient outcomes. ML algorithms have gained attention for their potential to enhance diabetes detection and prognosis by analyzing complex and nonlinear medical data (*22*). The following aims to provide a comprehensive overview of the ML approaches employed for diabetes detection and prognosis. By critically examining the existing research, we aim to identify the strengths and limitations of different techniques and highlight potential avenues for our proposal.

Ahmed et al. (*23*) developed an optimized ML-based classifier model for diagnosing diabetes using clinical data. Their approach included effective preprocessing techniques and achieved superior efficiency compared to existing methods, with an improvement in accuracy ranging from 2.71% to 13.13%. However, the generalizability of their model to different datasets and populations requires further investigation.

Hasan et al. (*18*) proposed a comprehensive architecture for diabetes prognosis, incorporating outlier exclusion, data normalization, and weighted ensembling of multiple ML models. Their suggested ensemble model achieved an Arear Under Curve (AUC) score of 95% on the Pima Indian dataset. However, the study’s limitation is that it focused only on the performance of a single dataset, limiting the assessment of generalizability.

Howlader et al. (*24*) applied ML strategies to identify Type 2 Diabetes (T2D) patients. They performed extensive feature selection and analysis using various classification algorithms, with the Generalized Boosted Regression (GBR) model achieving the best accuracy rate of 90.91%. However, the study’s scope was limited to the prediction of T2D and did not explore other types of diabetes or broader diabetes prognosis.

Deepajothi et al. (*25*) aimed to forecast diabetes in its initial phases by incorporating hereditary factors into a fuzzy classification model. Their suggested model achieved an accuracy rate of 83% for identifying Type 2 diabetes using the Pima Indian dataset. However, the study’s limitation is that it did not compare the performance of their models with other existing diabetes prognosis methods.

Rajagopal et al. (*26*) developed a modified combined approach of artificial neural networks (ANN) with genetic algorithms for diabetes detection. Their model achieved an accuracy rate of 80% on the Pima Indian dataset. However, the study did not explore the performance of other ML algorithms’ performance or evaluate their approach’s generalizability on different datasets.

Nuankaew et al. (*27*) proposed a unique predicting approach called Average Weighted Objective Distance (AWOD) for diabetes forecasting. Their technique achieved an accuracy rate of 93.22% on the Pima Indian dataset and 98.95% on the Mendeley dataset. However, the study did not compare the performance of AWOD with other existing diabetes prediction methods.

Wei et al. (*28*) developed a methodology to estimate the usefulness of ambient chemical exposure in diagnosing diabetes mellitus. Their ML model utilizing the least absolute shrinkage and selection operator (LASSO) regression achieved an AUC of 80% for diabetes detection. However, the study’s limitation is that it focused only on the prediction of diabetes and did not consider other aspects such as prognosis or subtype classification.

Sivaranjani et al. (*29*) used Support Vector Machine (SVM) and RF ML algorithms to predict the likelihood of developing diabetes-related disorders. The RF model achieved an accuracy of 83% after feature selection and Principal Component Analysis (PCA) dimensionality reduction. However, the study did not explore other classification algorithms’ performance or evaluate their approach’s generalizability on different datasets.

Ramesh et al. (*30*) introduced an end-to-end monitoring system for diabetes risk stratification and control. Their SVM model achieved an accuracy rate of 83.20% using the Pima Indian dataset. However, the study’s limitation is that it focused on risk stratification and did not extensively evaluate the performance of their model on other aspects such as diagnosis or prognosis.

Ravaut et al. (*31*) constructed an ML-based Gradient Boosting Decision Tree (GBDT) model for estimating adverse outcomes related to diabetes. Their model achieved an AUC statistic of 77.7% for predicting the three-year risk of developing diabetes complications. However, the study’s limitation is that it relied on organizational health data from a specific region, and the generalizability of their model to other populations requires further investigation.

Naz et al. (*32*) proposed a procedure for early diabetes estimation using various ML classifiers and the Pima dataset. Their deep learning (DL) approach achieved a success rate of 98.07% and retrieved viable properties. However, the study’s limitation is that it focused solely on DL-based methods and did not explore the performance of other ML algorithms.

Hassan et al. (*33*) provided a diabetes prognosis model based on ML algorithms, achieving accuracy rates of 94.5%, 96.5%, and 97.5% for Logistic Regression (LR), SVM, and RT, respectively. However, the study’s limitation is that it utilized a relatively small dataset of 250 variations, raising questions about the generalizability of its model to larger and more diverse datasets.

Gupta et al. (*34*) developed prognostic tools using DL and quantum ML (QML) approaches. Their DL classifier achieved an accuracy rate of 95%, while the QML classifier had 86% accuracy. However, the study’s limitation is that it focused on comparing DL and QML methods without considering other traditional ML algorithms.

Gupta et al. (*35*) introduced the application of Moth-Flame Optimization (MFO), a metaheuristic algorithm, for classifying diabetes data. They incorporated MFO to update feedforward neural network weights and conducted performance evaluations on the Wisconsin Hospital dataset, along with a comparative analysis of contemporary literature.

Majhi et al. (*36*) investigated and compared multiple ML approaches for early diabetes risk assessment and medical diagnosis enhancement. Their study employed two realworld datasets: a diabetic clinical dataset (DCA) from Assam, India, and the publicly available PIMA Indian diabetic dataset. Various classifiers were utilized, with logistic regression yielding the most promising results on PIMA, achieving an accuracy of 79.22%.

In contrast to these studies, our current work aims to address the limitations mentioned above. We propose an optimized data preprocessing pipeline, tackle imbalanced datasets, prevent overfitting using k-fold cross-validation, and conduct extensive experimental validation on diverse datasets.

It is undoubtedly challenging to predict diabetes in its early stages due to the complex interdependencies between numerous factors. Creating a medical prediction model that aids medical professionals in the prediction procedure is necessary. An accurate diabetes prognosis is crucial to prevent premature death. Therefore, achieving a greater accuracy rate with a lower error rate is required to forecast diabetes better. In this paper, we have adopted several machine-learning algorithms to predict diabetes and identify the best one based on clinical data related to diabetes to address these issues. To improve accuracy and achieve higher performance, it is necessary to preprocess the raw data to match the criteria of different classifiers. An efficient data preprocessing pipeline is provided to the learning algorithms for predictive modeling. To assess our proposal, extensive experimental analysis has been performed on four diabetes datasets, each with different attributes. Experimental results show that our proposal surpasses state-of-the-art research in predicting diabetes, with an average accuracy of 95.5%.

The paper makes the following contributions:

### Optimized data preprocessing pipeline

We develop a robust pipeline for preprocessing diabetes-related datasets. This includes handling missing values, outliers, label encoding, and normalization. Our preprocessing techniques improve dataset quality, leading to enhanced classifier performance.

### Addressing imbalanced datasets

We tackle the challenge of imbalanced data by implementing random oversampling techniques. This creates a balanced dataset, improving the performance of diabetes detection and prognosis models.

### Overfitting prevention

To prevent overfitting, we employ k-fold cross-validation during model training. This ensures that the models generalize well to unseen data, enhancing their reliability for diabetes prediction.

### Extensive experimental validation

Through extensive experiments on diverse diabetes datasets, our approach consistently outperforms existing methods. We demonstrate superior accuracy, precision, recall, and F1-score, validating its effectiveness for diabetes detection and prognosis.

These contributions advance the field of diabetes research by providing an optimized preprocessing pipeline, addressing dataset imbalance, preventing overfitting, and demonstrating superior performance through extensive experimentation.]

The hypothesis of this paper is described as follows:

### Hypothesis

This study proposes an optimized data preprocessing pipeline and addresses the challenge of imbalanced datasets in the context of diabetes detection and prognosis. By implementing robust preprocessing techniques, including handling missing values, outliers, label encoding, and normalization, and employing random oversampling, we hypothesize that the dataset quality will be enhanced, leading to improved classifier performance. Furthermore, by employing k-fold cross-validation to prevent overfitting and conducting extensive experimental validation, we expect to demonstrate our approach’s superiority in accuracy, precision, recall, and F1-score compared to existing methods.

These findings will validate the effectiveness of our proposed methodology for diabetes detection and prognosis, making a significant contribution to the field.

The rest of the paper is organized as follows. In section 2, we review the related literature for diabetes detection. Sections 3 and 4 describe our proposal and various ML algorithms adopted in predictive learning. Experimental results are presented in Section 5. Then in Section 6, we present the implementation details of the web-based application and Finally, in Section 7, we conclude the paper with the future enhancement of this study.

### Proposed Methodology

In this section, we have described our proposed methodology along with the different machine algorithms adopted in the system. Firstly, we explain the working principle of the proposal. Then, we briefly describe the ML models.

Figure 1 shows the basic workflow of the proposed approach. The proposal has four significant parts: data collection, data preprocessing, handling imbalanced class problems, splitting datasets using k-fold cross-validation and applying ML algorithms to train and test the models, and evaluating the performance.

**Figure 1.**
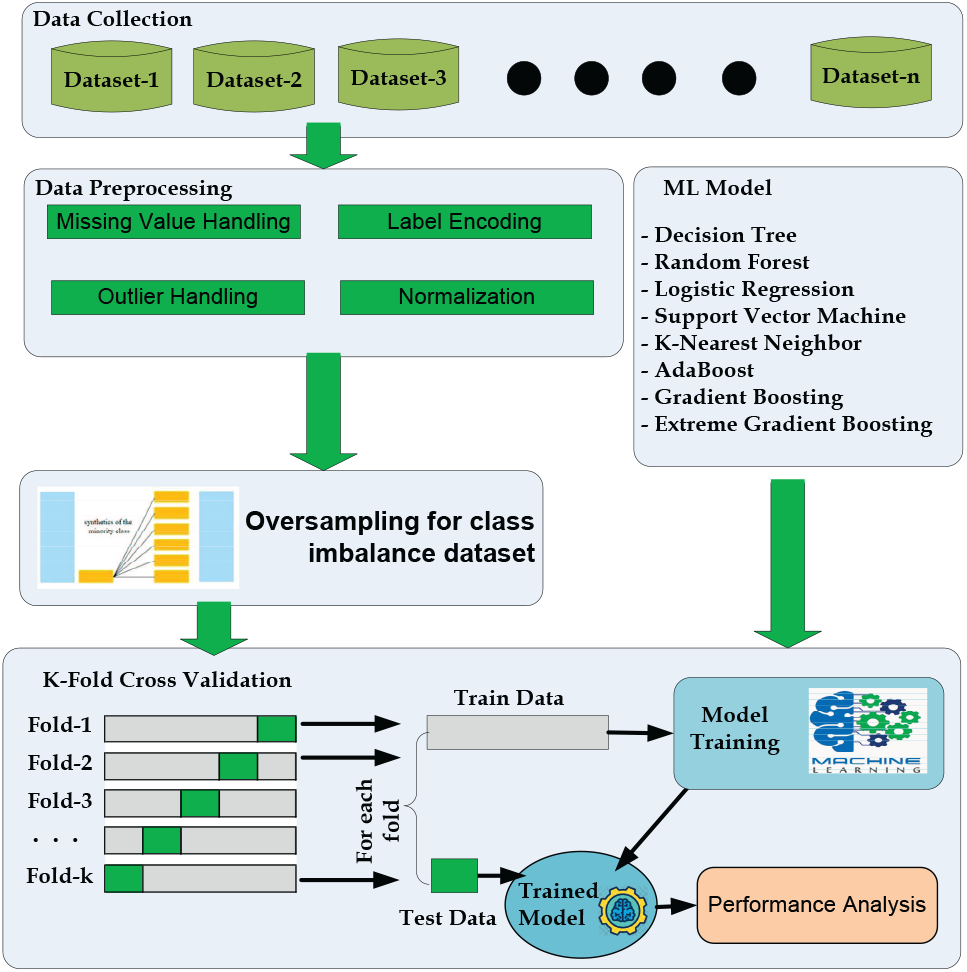
The workflow diagram for Diabetes prediction.

The steps performed in our study are as follows:

#### Data collection

We gathered the necessary data from publicly available sources. Data preprocessing: After data collection, we conducted data preprocessing to prepare the dataset for model training and evaluation. This includes cleaning the data, handling missing values, and performing outlier handling, label encoding, and data normalization.

#### Handling imbalanced class problems

To address any class imbalance issues in the dataset, we employed techniques such as oversampling, undersampling, or synthetic data generation to balance the classes. This step aims to prevent biases in the model due to imbalanced data.

#### Splitting datasets using k-fold cross-validation

We utilized k-fold cross-validation to split the dataset into training and testing folds. The preprocessing steps, including data balancing, were applied within each fold. This ensures that the data preprocessing steps are performed independently for each fold and prevents any information leakage between the training and testing sets.

#### ML model building and evaluation

Within each fold, we built ML models using the training data and evaluated their performance on the corresponding testing data. This process was repeated for each fold, resulting in multiple model evaluations.

By following this pipeline, we aimed to ensure a robust evaluation of our proposed method for diabetes prediction. In our proposed methodology shown in Figure 1, the “Data Preprocessing” box is placed outside the K-fold crossvalidation loop in this proposal for the following reasons:

#### Preventing Data Leakage

By performing data preprocessing separately for each fold, outside the crossvalidation loop, information leakage from the test set to the training set is avoided. This ensures unbiased evaluation and a more accurate assessment of the model’s generalization capabilities.

#### Realistic Evaluation

Applying data preprocessing techniques outside the cross-validation loop mimics real-world scenarios where models encounter unseen, preprocessed data during deployment. This approach provides a realistic evaluation of the model’s effectiveness and its performance on truly unseen data.

#### Efficiency

Placing data preprocessing outside the cross-validation loop improves computational efficiency. Preprocessing is performed once on the entire dataset before cross-validation, reducing redundant computations within each fold and speeding up the overall evaluation process.

It’s important to consider the specific requirements of the study and the nature of the preprocessing techniques used when deciding the placement of data preprocessing and cross-validation to ensure unbiased and reliable model evaluation.

### Data Collection

To ensure the robustness of our model, we gathered data from four distinct datasets, each containing different variables related to diabetes. These datasets were collected from various sources, including demographic data, diabetes statistics, and health characteristics obtained from individuals across different countries and healthcare institutions. The first dataset used in our study is the Pima Indian Diabetes Dataset (*37*), which is widely recognized as a valuable resource for evaluating ML algorithms in predicting diabetes within the general population. Dataset 2 (*38*), referred to as the Austin Public Health Diabetes Self-Management Education Participant Demographics 2015-2017, comprises demographic information collected from participants in the diabetes self-management education program conducted by Austin Public Health. For Dataset 3 (*39*), a survey was conducted to gather data from 950 records, including 19 attributes that have been identified as having a measurable influence on diabetes. Lastly, Dataset 4 (*40*) was collected from the Iraqi society, as well as from the laboratory of Medical City Hospital and the Specialized Center for Endocrinology and Diabetes at Al-Kindy Teaching Hospital. By utilizing these diverse datasets, we aim to enhance the generalizability and applicability of our proposed model for diabetes prediction and prognosis.

### Data Analysis and Data Preprocessing

Data preprocessing is the process of preparing original data for ML. It is indeed the most important step in the process of developing an ML model. It is a necessary step for ML algorithms to improve the model’s accuracy and efficiency.

Data preprocessing covers data preparation, which includes data integration, cleansing, normalization, and transformation, as well as data reduction activities like feature selection, instance selection, and discretization.

We have conducted some data analysis after collecting the data. Then, we have done preprocessing tasks including outliers removal and dealing with missing values, data normalization, encoding, etc.

#### 1. Outliers Handling

A dataset may contain extreme values that are beyond the acceptable limits and dissimilar to the rest of the data. This kind of data may reduce the performance of the ML algorithm. Any value ≤ *Q*_1_ −1.5 ×*IQR* or ≥ *Q*_3_ −1.5 × *IQR* is considered as outlier. Identifying and handling outliers can be expressed as the following algorithm 1 using first quartile *Q*_1_, third quartile *Q*_3_ and Interquartile Range (IQR),:

#### 2. Missing value handling

One of the most difficult tasks for analysts is dealing with missing values, because making the proper decision on how to deal with them leads to more robust data models. We can handle missing values in various ways, like ignoring the row, imputation of the missing values with data means, median or mode of the observation, imputation with different ML algorithms or prediction using regression, etc. To increase the model performance, the mean value of the respective attribute has been imputed to manage the missing values, which are crucial for diabetes prediction and can be calculated in Equ. 1 as follows:

##### Algorithm 1 Handling Outlier Problem

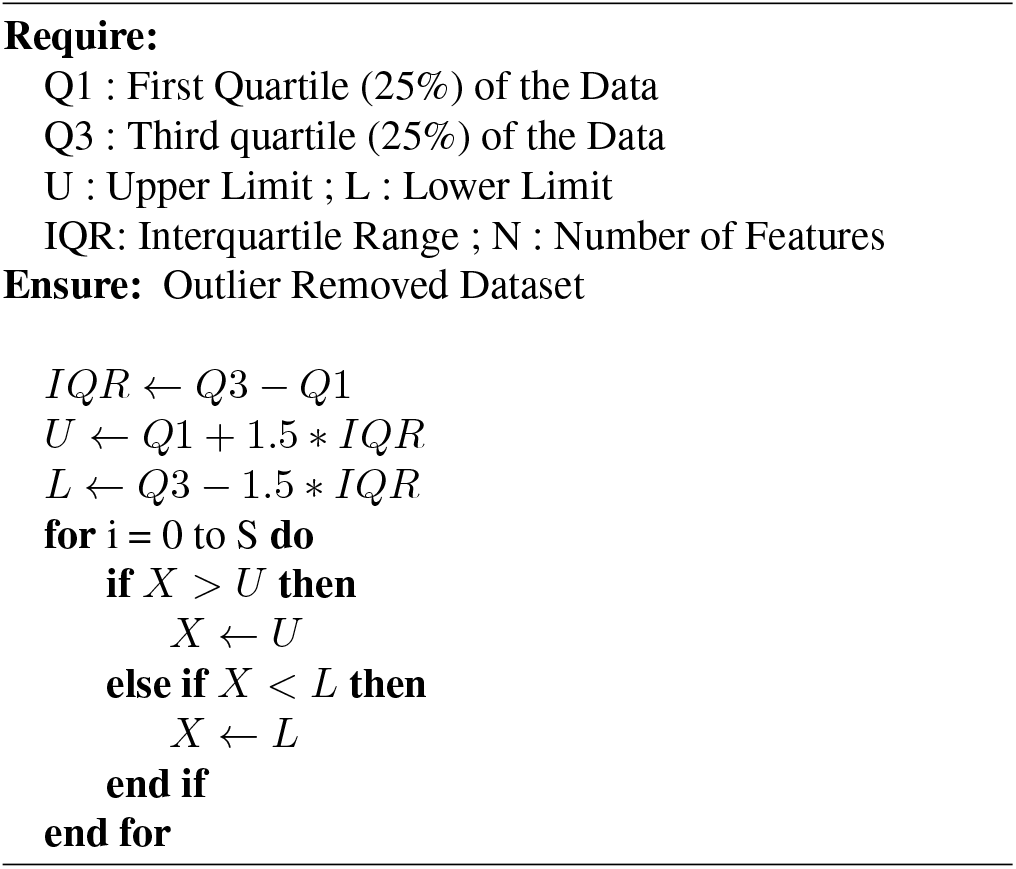

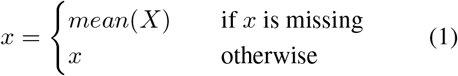

here, the value of instance *x* is imputed with the attribute mean as denoted as *mean*(*X*).

#### 3. Label Encoding

All the data entered in the ML algorithm must be numerical. However, in most cases, the dataset may contain categorical data rather than numerical values. So, if the dataset contains any categorical attribute, it must be needed to convert to numeric values before fitting and evaluating an ML model. Label encoding is the process of transforming labels of text/categorical values into a numerical format which is understandable by the ML algorithms. For example, we can convert the categorical values of *Gender* status ‘Male’ to ‘1’ and ‘female’ to ‘0’.

#### 4. Standardization-

When features of an input data set have considerable variations between their ranges or when they are collected or measured in different measurement units, standardization becomes necessary. Differences hamper the performance results for ML models in the range of initial features. So normalization or standardization can solve this issue and improve the prediction quality. We have used standardization to rascal the values of any attribute for better accuracy of the classification model using the following Equ. 2:

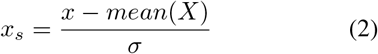

here, *x*_*s*_, *mean*(*X*) and *σ* represent the standardization value, mean value, and standard deviation of the attribute *X*.

### Class balancing using oversampling

There is a class imbalance when observation in one class exceeds observation in other classes. This may cause poor performance as the ML algorithm may ignore the minority class. To deal with this problem, we apply the random oversampling method to balance the dataset. The benefit of oversampling is that no information from the original training set is lost because all members of the minority and majority classes are kept, and it also significantly increases the size of the training set.

### ML Algorithms

We have used 7 different ML classifiers to train the model, including DT, NB, KNN, LR, XGB, and SVM, and predict diabetes. Depending on several performance metrics, the performance of each classifier has been analyzed. The following subsection will describe the ML algorithms used for the predictive model.

#### Naive Bayes (NB)

NB is based on Bayes’ Theorem (*41*). It assumes that the value of one feature in a class is independent of the presence of any other feature. Despite its simplicity, Naive Bayes often outperforms more sophisticated classification methods (*23*).

#### Decision Tree (DT)

DT is a supervised classifier commonly used for solving classification problems (*42*). It effectively captures decision-making information from the dataset. Internal nodes represent features, and leaf nodes represent outcomes.

#### Random Forest (RF)

RF is an ensemble classifier that trains multiple decision trees (*43*). The final classification is based on the majority vote of all trees. Increasing the number of trees generally improves performance.

#### Logistic Regression (LR)

LR is a supervised classifier used for binary classification tasks based on event probabilities (*17*). It assumes linear separability of data.

#### K-Nearest Neighbor (KNN)

KNN is a supervised learning method for classification (*44*). It assigns the new case to the class nearest to it in the training dataset.

#### Gradient Boosting (GB)

GB predicts continuous or categorical target variables by iteratively improving models (*45*). It corrects errors from previous models, enhancing overall performance.

#### Extreme Gradient Boosting (XGB)

XGB is a gradient-boosted decision tree implementation known for its speed and efficiency (*43*).

#### Adaptive Boosting (AdaBoost)

AdaBoost is an ensemble learning technique that improves the performance of ML algorithms, often using decision trees with only one split (*46*).

#### Support Vector Machine (SVM)

SVM efficiently separates datasets by finding optimal decision boundaries or hyperplanes in n-dimensional space, maximizing the margin between support vectors (*47*).

### K-fold cross validation

Cross-validation is a resampling technique for evaluating ML models. It guarantees that every observation in the dataset can be selected in the training and test data. It’s one of the best tactics if we have limited raw data. A badly chosen value for k could result in an inaccurate illustration of the model’s skill, such as accuracy with a high variance. A k=10 means the kfold process divides the whole dataset five or ten times for evaluating the model. For our experiments, we use k=10 fold validation and take the performance average for the result analysis.

## Results

We have conducted our proposal using various preprocessing techniques as well as utilizing several ML algorithms to analyze and find the best model to use in the prediction of diabetes for clinical purposes. We have tested our proposal with 4 different datasets and each of them contains different types and numbers of attributes to prove that our preprocessing techniques are more efficient than the normal preprocessing process as well as other existing research works.

### Experimental Setup

All the experiments were conducted on a computer having an Intel Core i7 processor with a 4 GB graphics card, 16GB RAM, and a 64-bit Windows operating system running at 1.80 GHz using the Python programming language. The dataset is shuffled and divided into 10 folds at random, with one fold used for the testing and the others used for training each ML model. Then, the resultant average value of any performance results has been taken to assess them.

### Performance Metrics

The prediction of any ML algorithm could have four distinct results depending on the confusion matrix as indicated in the Table 1: true positive (TP), true negative(TN), false positive(FP), and false-negative(FN).

**Table 1.** Confusion Matrix.

| Predicted Results | Actual positive | Actual negative |
| --- | --- | --- |
| Yes | TP | FP |
| No | FN | TN |

Then we consider the following metrics to analyze the proposal:

- Accuracy measures the overall correctness of the model’s predictions. It is calculated as the ratio of the number of correct predictions (true positives and true negatives) to the total number of predictions made (Equ. 3).

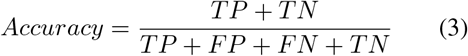
- Precision quantifies the ability of the model to avoid false positives. It is calculated as the ratio of true positives to the sum of true positives and false positives (Equ. 4).

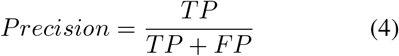
- Recall measures the model’s ability to capture all positive instances. It is calculated as the ratio of true positives to the sum of true positives and false negatives (Equ. 5).

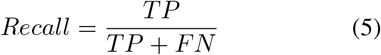
- The F1-score is the harmonic mean of precision and recall, providing a balance between the two metrics. It is calculated as twice the product of precision and recall divided by the sum of precision and recall (Equ. 6).

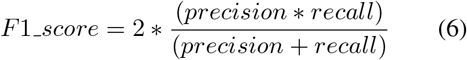
- Mean Absolute Error (MAE) measures the average absolute difference between predicted values and actual values. It is calculated as the sum of absolute differences divided by the total number of instances (Equ. 7).

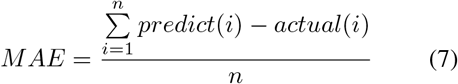
- Mean Squared Error (MSE) measures the average of the squares of the errors or deviations. It is calculated as the sum of squared differences divided by the total number of instances (Equ. 8).

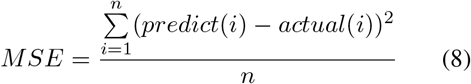
- Root Mean Squared Error (RMSE) is the square root of the MSE and represents the average magnitude of the error. It provides a measure of how spread out the errors are (Equ. 9).

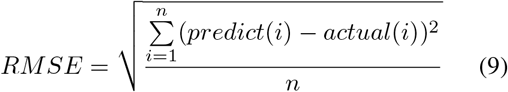
- The Area under Receiver Operative Curve (ROC) (AUROC) quantifies a binary classifier’s ability to distinguish between classes. It ranges from 0.5 to 1, where higher values signify better performance. In medical settings, AUROC assesses a test’s accuracy in identifying patients with a condition while minimizing misclassifications. It offers a concise measure of the model’s overall discriminative power, crucial for evaluating diagnostic tools in clinical practice.

### Hyperparameters

Hyperparameter selection is an important aspect of ML model development, and we acknowledge that our previous submission lacked information on the hyperparameter optimization process. Here, we provided an overview of the hyperparameter values used for each ML model in Table 2.

**Table 2.**
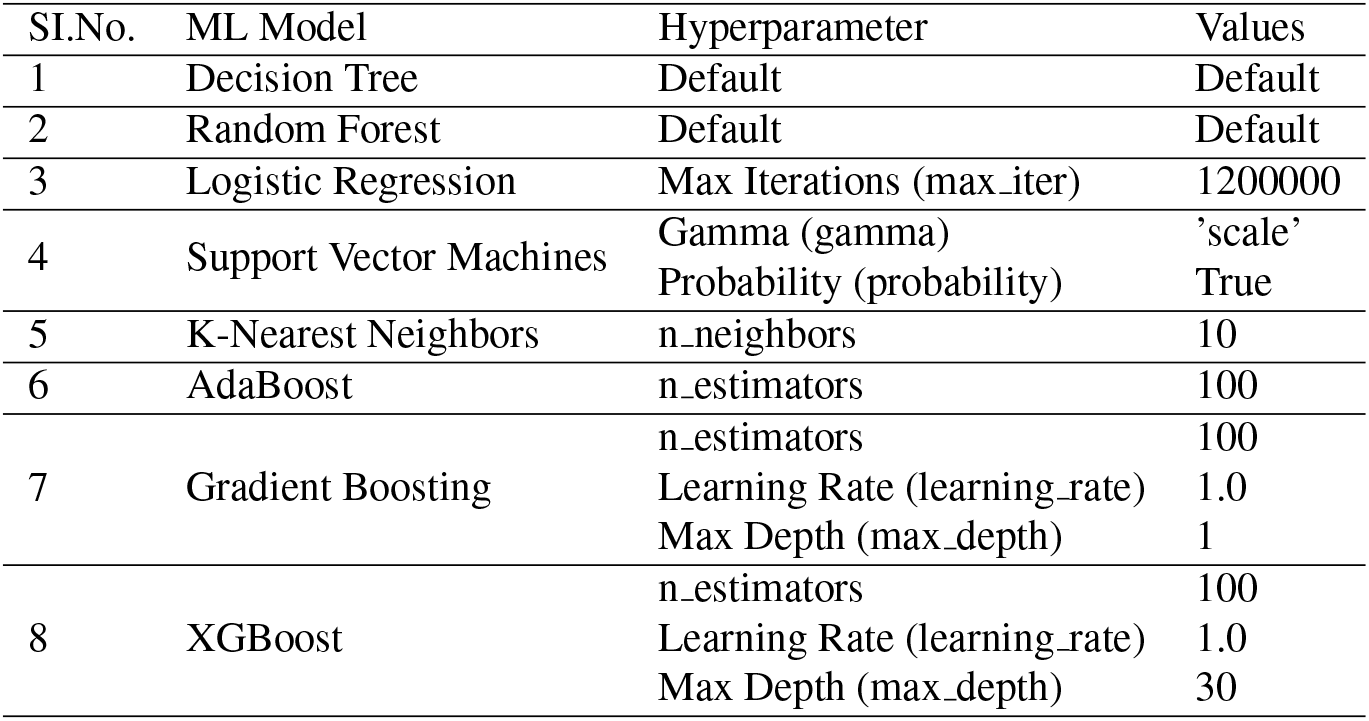
Hyperparameter tuning values for ML models.

In the case of the Decision Tree (DT) and Random Forest (RF) models, we used the default hyperparameter values as provided by the respective implementations. For Logistic Regression (LR), we set the maximum number of iterations (max iter) to 1,200,000. For Support Vector Machines (SVM), we used a gamma value of ‘scale’ and enabled probability estimation by setting the probability parameter to True. The K-Nearest Neighbors (KNN) model utilized a value of 10 for the number of neighbors (n neighbors). For AdaBoost (ADB), we set the number of estimators (n estimators) to 100. In the case of Gradient Boosting (GBC), we used 100 estimators, a learning rate of 1.0, and a maximum depth of 1. Lastly, the XGBoost (XGB) model employed 100 estimators, a learning rate of 1.0, and a maximum depth of 30.

We acknowledge the importance of hyperparameter optimization in improving model performance. In our initial submission, we unintentionally omitted the details of our hyperparameter selection process. We apologize for this oversight and recognize the significance of following best practices in ML.

### Results of Dataset -1

The Pima Indian Diabetes Dataset is one of the most useful datasets for testing ML algorithms for predicting diabetes in the general population (*37*). This dataset was provided by the National Institute of Diabetes and Digestive and Kidney Diseases and is used to determine whether a patient has diabetes based on diagnostic measures such as pregnancy, glucose level, blood pressure, skin thickness, diabetes pedigree function, insulin, body mass index (BMI), and age. The attributes of the Pima Indian dataset are listed below:

1. Pregnancies:Number of occurrences of pregnancy
2. Glucose:In a glucose tolerance measure, the plasma glucose concentration after 2 hours
3. Skin Thickness:The thickness of the skin folds on the triceps (mm)
4. Insulin:Serum insulin (mu U/ml) after 2 hours
5. BMI:Body mass index
6. Age:Age of the person in years
7. Outcome:Class variable as a result (0 or 1)

The boxplot in Figure 2(a) shows that the dataset contains outliers, whereas Figure 2(b) shows clean data after applying the preprocessing algorithm. In the boxplot, different features have multiple outliers data which is indicated by multiple diamond signs beside each feature and after handling the outlier the boxplot looks like no diamond signs on each feature which proves no outlier existed.

**Figure 2.**
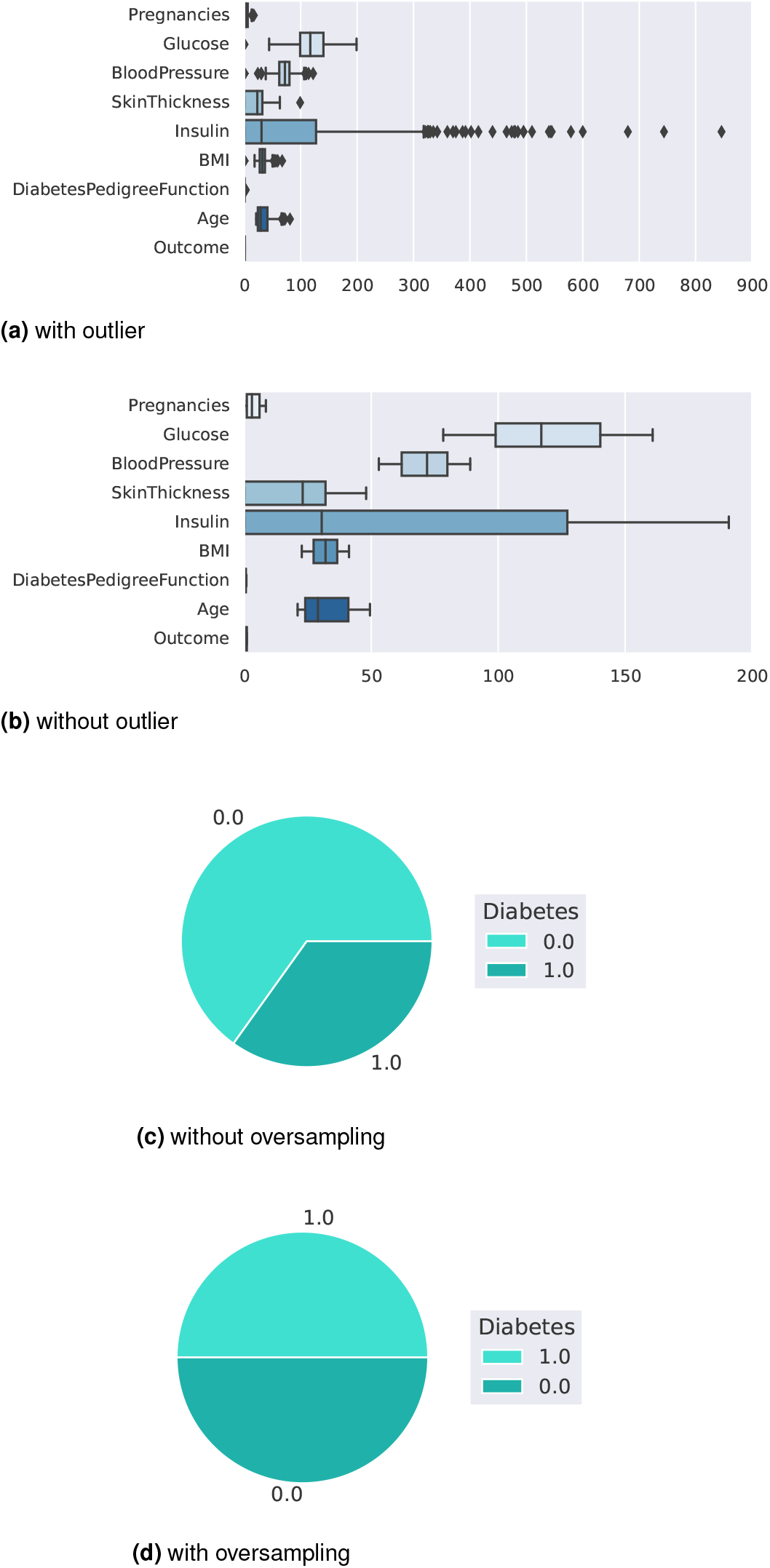
Before and after outlier removal and oversampling results

Figure 2(d) depicts the balanced dataset distribution of the original imbalance dataset 2(c), where label 0 represents no diabetes and 1 represents diabetes. In the pie chart, we can see that it contains more portion of 1 than 0 and after random oversampling, we can see that we have the same portion of labels 0 and 1, ensuring data is balanced now.

Figures 3(a, b) represent the accuracy and MSE of our experiments for dataset-1. The accuracy comparison before and after applying the proposal. The accuracy resultss of DT, RF, LR, SVM, KNN, AdB, GB and XGB are 77.27%, 85.53%, 78.95%, 78.95%, 80.26%, 80.26%, 81.58% and 83% respectively. We found that, depending on the ML algorithms, accuracy performance increases from 4.95% to 12.15%. On the other hand, the MSE values of the algorithm reduced significantly, 5.27%, 12.15%, 5.95%, 4.95%, 8.26%, 6.23%, 8.85% and 10.27% for DT, RF, LR, SVM, KNN, AdB, GB and XGB respectively.

**Figure 3.**
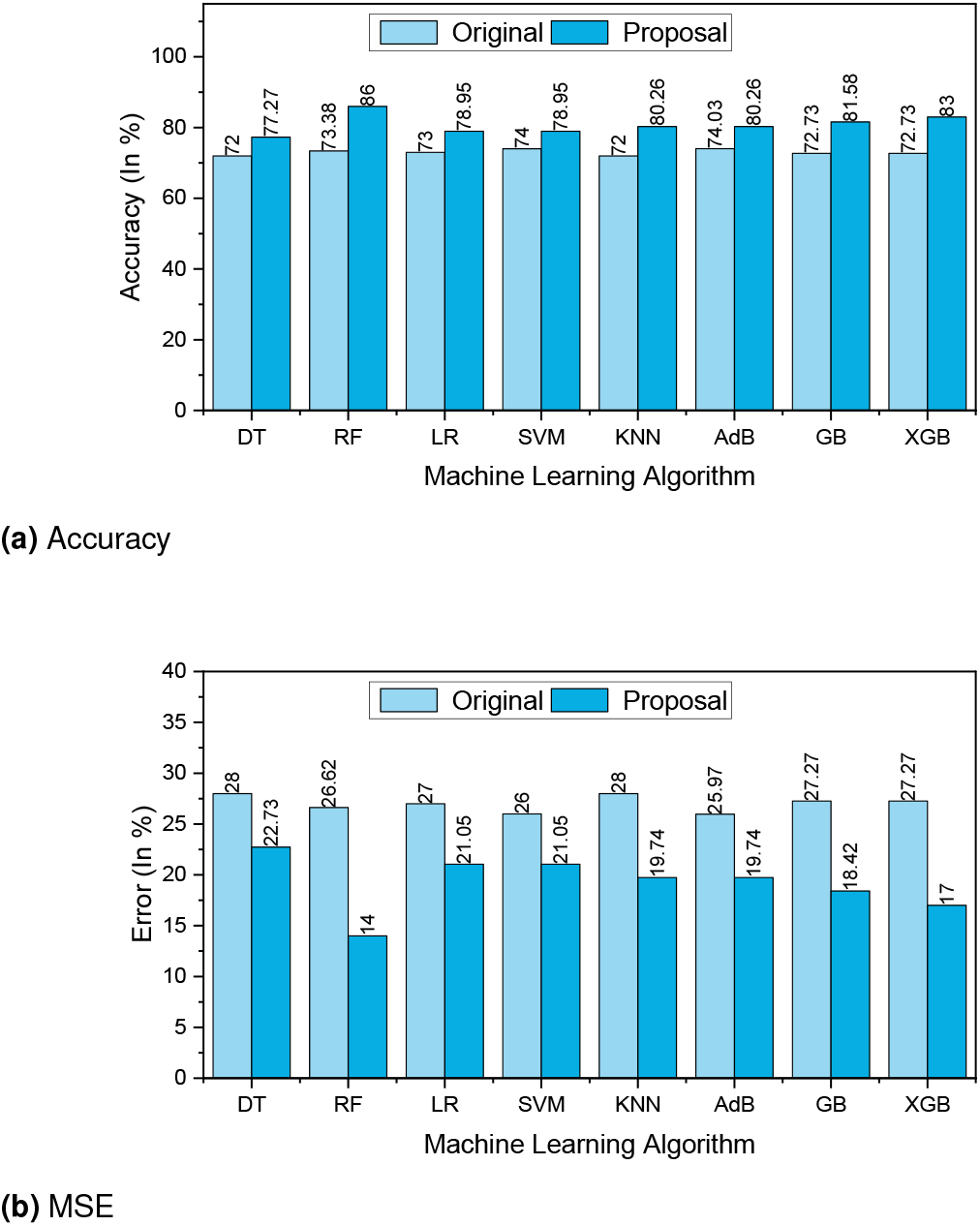
The performance Results of Dataset-1.

**Figure 4.**
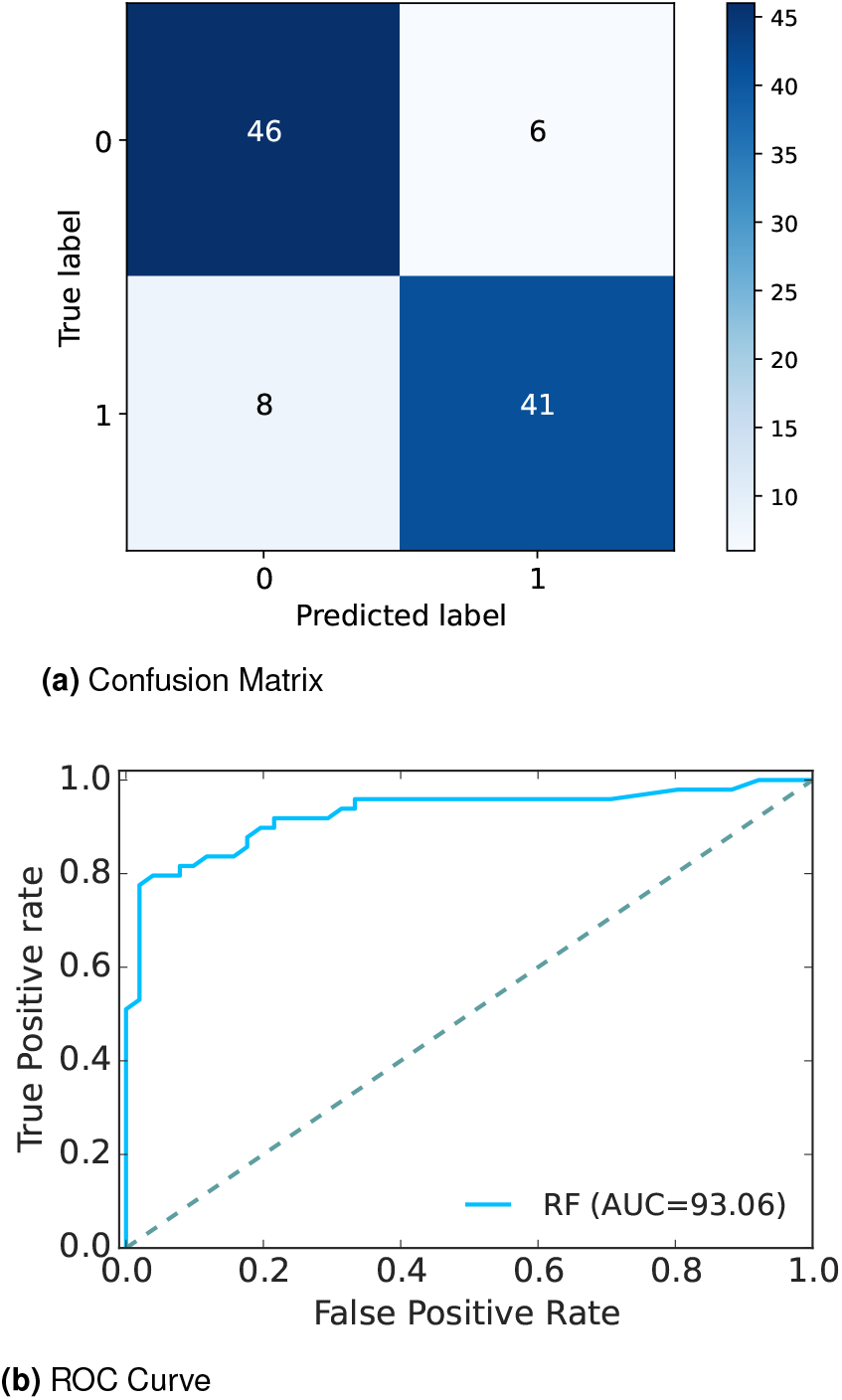
Confusion Matrix and ROC Curve of Dataset-1

Table 3 summarizes the other performance metrics. We found that the proposal can improve the precision values from 3.03% to 15.98% for any ML approach. An efficient data preprocessing and data balancing can improve the data quality; hence ML algorithms can accurately classify the test data. We found similar results for recall; the values increased from 0.41% to 11.94%. The F1-score also improved as expected, from 1.1% to 12.25%. On the other hand, the table also indicates that due to the high performance of the proposal, the values of MAE and RMSE are reduced significantly. It is observed that MAE values reduced from 4.95% for SVM to 12.15% for RF. Similarly, RMSE reduced greatly from 5.11% to 13.56%.

**Table 3.**
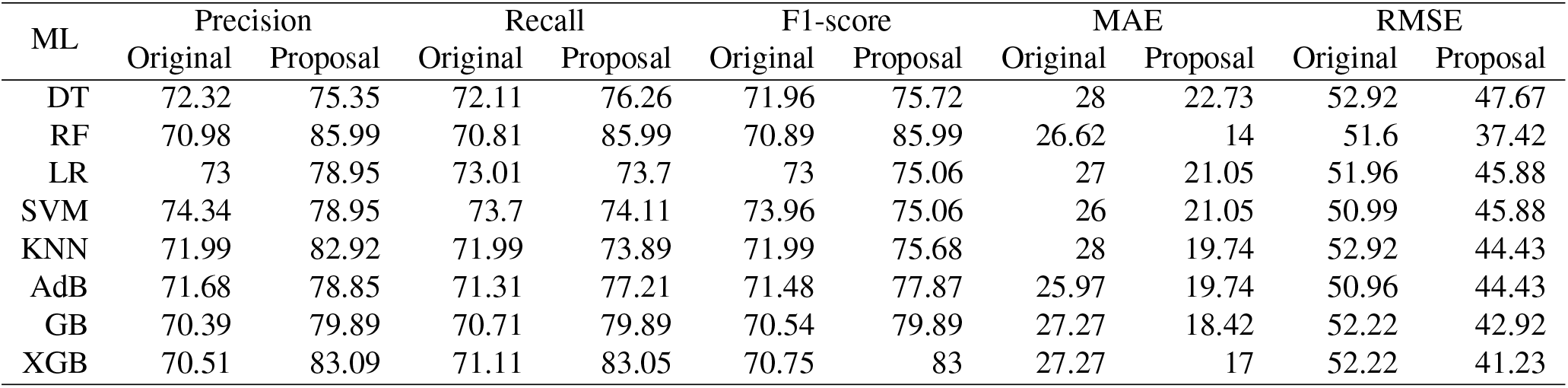
Performance result of dataset-01 (Pima Indian dataset)

The extensive analysis of the experiments shows that RF outperforms others as it is an ensemble classifier that works by training a large number of decision trees, leading to higher accuracy with good reliable predictions than other single algorithms. So, we consider this model as our proposed model.

Graph 4(a) depicts the confusion matrix and 4(b) shows the ROC Curve for our proposed model. In the confusion matrix, the large number of TP and TN than FP and FN is a very crucial point to be a better prediction model for ML. In graph 4(a), we can see that the TP and FN rates are high and FN and FP rates are very low which gives a better sign of diabetes prediction. The TP, TN, FP, and FN rates are 45%, 41%, 8%, 6% respectively. On the other hand, in the ROC, the value of AUC which is near 1 is the best model for ML. In graph 4(b), we can see that the AUC value is .93 (93.07%) which means the positive and negative labels are mostly segregated, and the model is effective.

After analyzing multiple performance indicators, we can find that among all the ML algorithms RF outperforms others with an accuracy rate of 86%, 14% MSE rate, and 8% FP as well as 6% FN rates.

### Results of Dataset -2

Dataset 2 (*38*) namely, Austin public Health diabetes self-management education participant demographics 2015-2017, contains demographic information collected from Austin public health, Austin. Among the 1,688 rows, 25 columns; various important attributes are considered for our experiment including diabetes status, health indicators, health behaviors including Race/Ethnicity Diabetes Status, Heart Disease, High Blood Pressure, Tobacco Use, Previous Diabetes Education, Diabetes Knowledge, Fruits and Vegetable Consumption, Sugar-Sweetened Beverage Consumption, etc. The attributes of the Austin Public Health dataset are listed below:

1. Age:Age in year
2. Gender:Gender of the patient
3. Race/Ethnicity:Race/ethnicity of participant
4. Heart Disease:Heart disease diagnosis (yes/no)
5. High Blood Pressure:High blood pressure diagnosis (yes/no)
6. Tobacco Use:Tobacco user (yes/no)
7. Previous Diabetes Education:Previous diabetes education reported by participant (yes/no)
8. Diabetes Knowledge:Self-reported knowledge of diabetes (poor/fair/good)
9. Fruits and Vegetable Consumption:Fruits and/or vegetables eaten each week
10. Sugar-Sweetened Beverage Consumption:Sugar-sweetened beverages consumed each week
11. Food Measurement:Number of times food was measured each week
12. Carbohydrate Counting:Number of times carbohydrates were counted each week
13. Exercise:Number of days participant exercised each week
14. Diabetes Status:Diabetes status (yes/no) of participant

The boxplot in Figure 5(a) shows that the dataset contains outliers, whereas Figure 5(b) shows clean data after applying the preprocessing algorithm. In the boxplot, different features have multiple outliers data which is indicated by multiple diamond signs beside each feature and after handling the outlier the boxplot looks like no diamond signs on each feature which proves no outlier existed.

**Figure 5.**
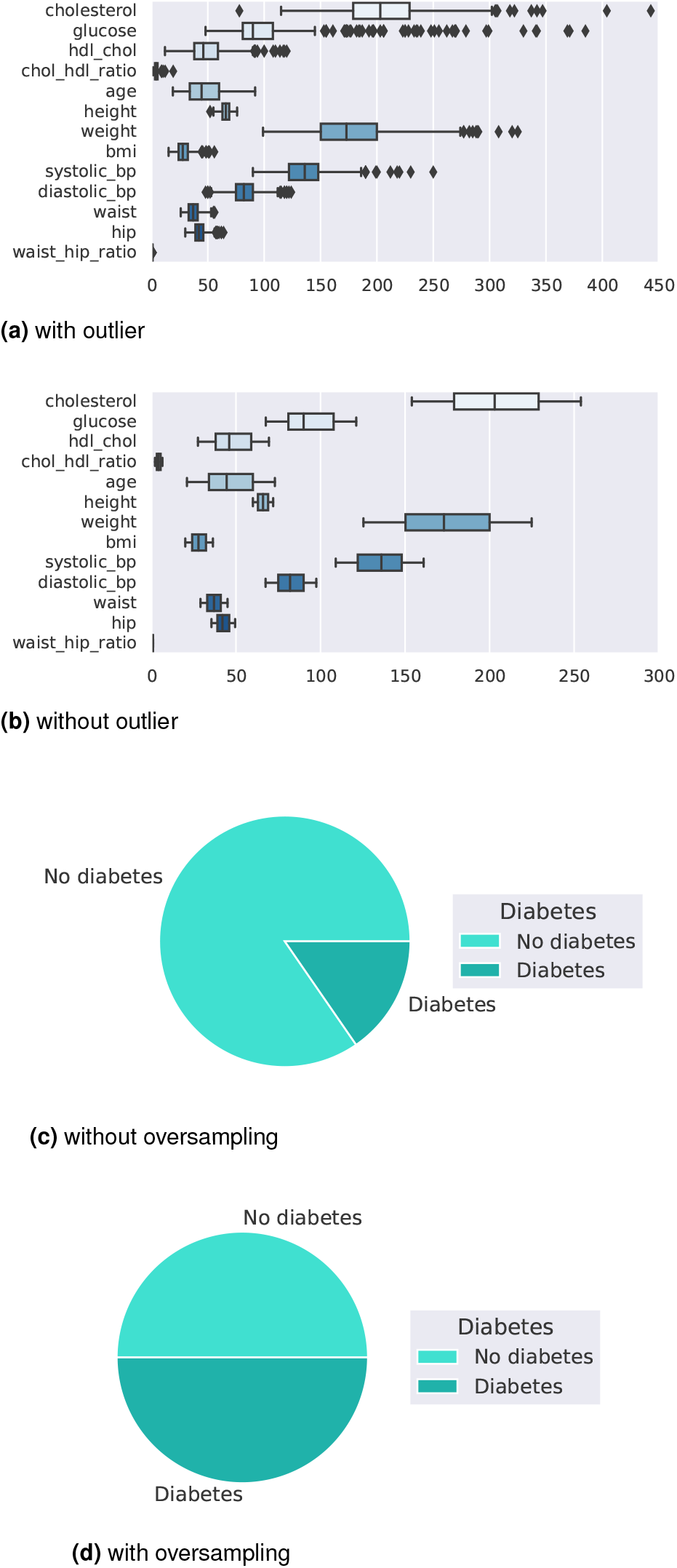
Before and after outlier removal and oversampling results

Figure 5(d) depicts the balanced dataset distribution of the original imbalance dataset Figure 5(c). where the label ‘No diabetes’ represents no diabetes and ‘Diabetes’ represents diabetes. In the pie chart, we can see that it contains more portion of ‘No diabetes’ than ‘Diabetes’ and after random oversampling, we can see that we have an equal portion of the labels ‘No diabetes’ and ‘Diabetes’ which ensures data is balanced now.

Figures 6(a, b) represent the accuracy and MSE of our experiments for dataset-2. The accuracy comparison before and after applying the proposal. The accuracy results of DT, RF, LR, SVM, KNN, AdB, GB and XGB are 95.45%, 98.48%, 83.33%, 93.94%, 87.88%, 96.97%, 93.94% and 93.94%, respectively. We found that, depending on the ML algorithms, accuracy performance increases from 0% to 8.74%. On the other hand, the MSE values of the algorithm reduced significantly, 5.71%, 8.74%, 0%, 5.48%, 1.52%, 5.94%, 4.2% and 1.63% for DT, RF, LR, SVM, KNN, AdB, GB and XGB respectively.

**Figure 6.**
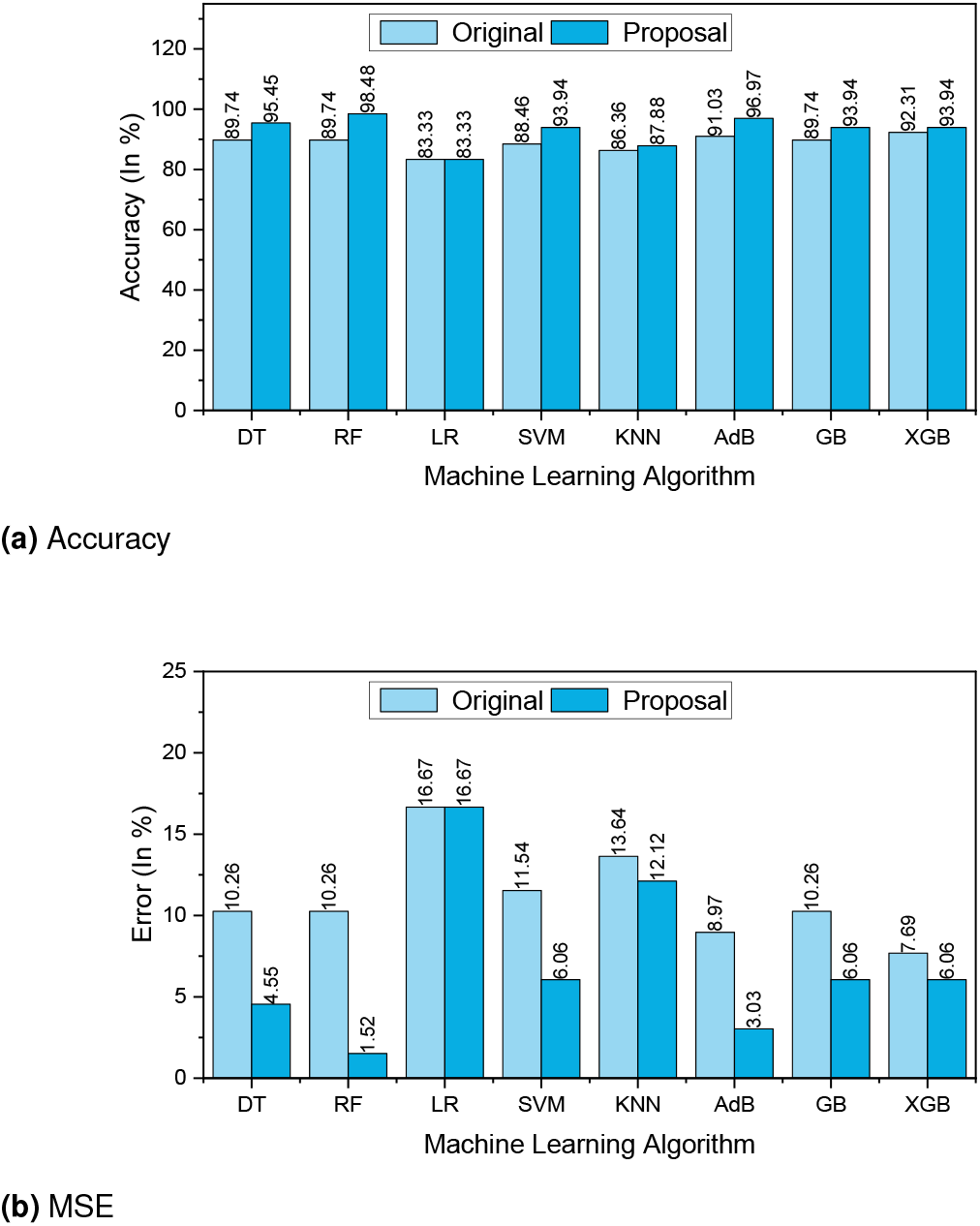
The performance Results of Dataset-2

**Figure 7.**
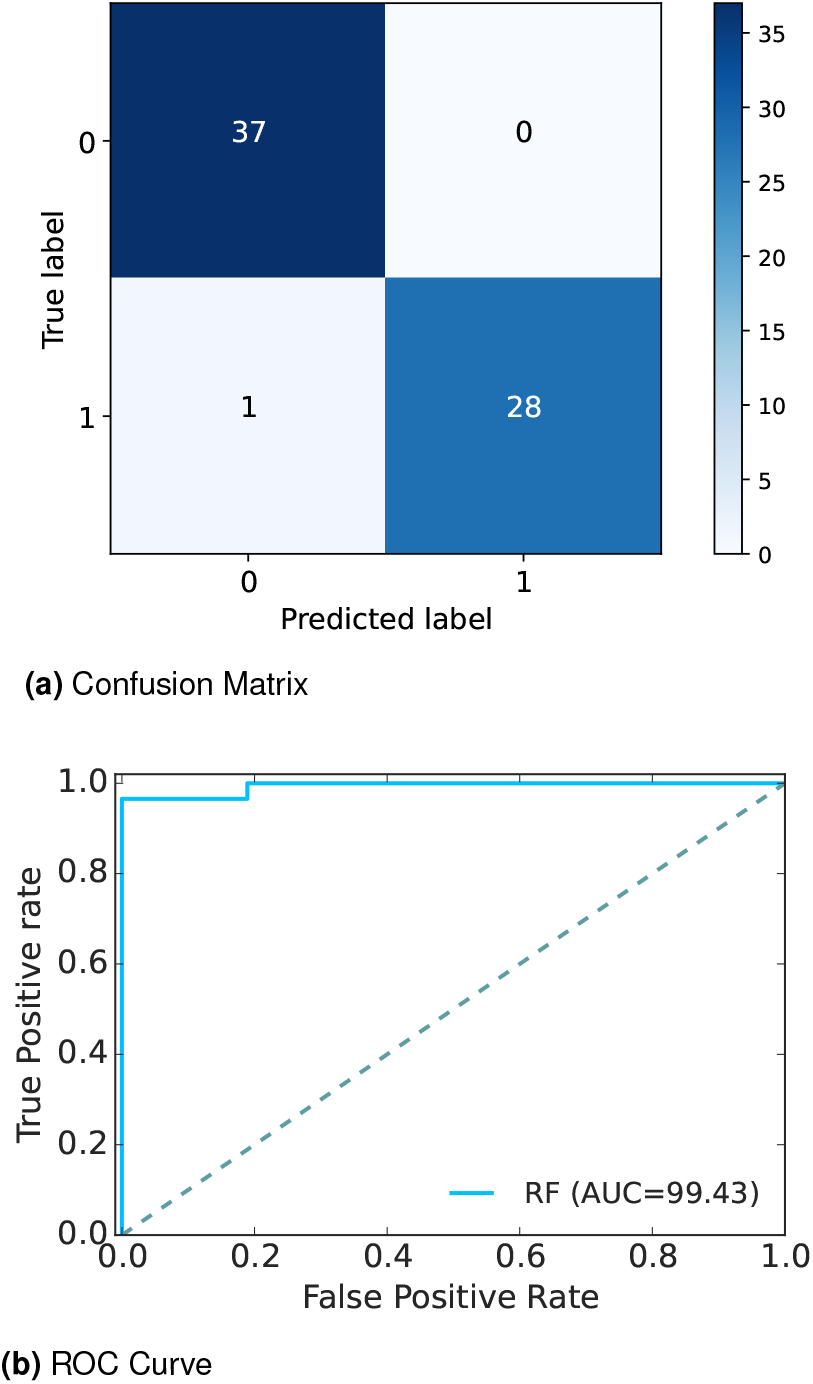
Confusion Matrix and ROC Curve of Dataset-2

The other performance metrics are summarized in Table 4. We found that, for any ML approach, the proposal can improve the precision values from 0% to 14.41%. An efficient data preprocessing and data balancing can improve the data quality, hence ML algorithms can accurately classify the test data. We found similar results for recall, the values increase from 8.27% to 20.57%. The F1 score also improved as expected from 0.47% to 14.19%. On the other hand, the table also indicates that due to the high performance of the proposal, the values of MAE and RMSE are reduced significantly. It is observed that MAE values reduced by 8.74% for RF to 0% for LR. Similarly, RMSE reduced greatly from 19.72% to 0%.

**Table 4.** Performance result of dataset-02 (Austin dataset)

| ML | Precision |  | Recall |  | F1-score |  | MAE |  | RMSE |  |
| --- | --- | --- | --- | --- | --- | --- | --- | --- | --- | --- |
|  | Original | Proposal | Original | Proposal | Original | Proposal | Original | Proposal | Original | Proposal |
| DT | 84.27 | 95.28 | 75 | 95.57 | 82.67 | 95.4 | 10.26 | 4.55 | 32.03 | 21.32 |
| RF | 84.27 | 98.68 | 84.27 | 98.28 | 84.27 | 98.46 | 10.26 | 1.52 | 32.03 | 12.31 |
| LR | 83.06 | 83.06 | 75 | 83.27 | 82.67 | 83.14 | 16.67 | 16.67 | 40.82 | 40.82 |
| SVM | 83.85 | 94.3 | 75 | 93.48 | 80.94 | 93.8 | 11.54 | 6.06 | 33.97 | 24.62 |
| KNN | 86.11 | 87.65 | 75 | 88.07 | 82.67 | 87.78 | 13.64 | 12.12 | 36.93 | 34.82 |
| AdB | 85.78 | 96.92 | 87.4 | 96.92 | 86.55 | 96.92 | 8.97 | 3.03 | 29.96 | 17.41 |
| GB | 84.27 | 93.73 | 84.27 | 94.22 | 84.27 | 93.89 | 10.26 | 6.06 | 32.03 | 24.62 |
| XGB | 87.22 | 93.73 | 75 | 94.22 | 82.67 | 93.89 | 7.69 | 6.06 | 27.74 | 24.62 |

The extensive analysis of the experiments shows that RF outperforms others so we consider this model as our proposed model. Graph 7(a) depicts the confusion matrix and 7(b) shows the ROC Curve for our proposed model. In the confusion matrix, the large number of TP and TN than FP and FN is a very crucial point to be a better prediction model for ML. In graph 7(a), we can see that the TP and FN rate are high and FN and FP rates are very low which gives a better sign of diabetes prediction. The TP, TN, FP and FN rates are 56.06%, 42.42%, 1.52%, 0% respectively. On the other hand, in the ROC Curve the value of AUC which is near 1 is the best model for ML. In graph 7(b), we can see that the AUC value is 0.99 (99.35%) which means the positive and negative labels are mostly segregated, and the model is effective.

After analyzing multiple performance indicators, we can find that among all the ML algorithms RF outperforms others with an accuracy rate of 98.48%, 0% MSE rate, and 1.52% FP as well as 0% FN rates.

### Results of Dataset -3

Diabetes-3 (*39*) conducted a survey and collected a dataset containing 950 records and 19 attributes that have a measurable influence on diabetes such as Family Diabetes history, Blood Pressure, Exercise, BMI, Smoking level, Alcohol consumption, Sleeping hours, Food habits, Pregnancy, Urination frequency, Stress level, etc. The attributes of the survey dataset are listed below:

1. Age:Age in Year
2. Gender:Gender of the participant
3. Family Diabetes:Family history with diabetes
4. highBP:Diagnosed with high blood pressure
5. PhysicallyActive:Walk/run/physically active
6. BMI:Body Mass Index
7. Smoking:Smoking
8. Alcohol:Alcohol consumption
9. Sleep:Hours of sleep
10. SoundSleep:Hours of sound sleep
11. RegularMedicine:Regular intake of medicine
12. JunkFood:Junk food consumption
13. Stress:Not at all, Sometimes, Often, always
14. BPLevel:Blood pressure level
15. Pregancies:Number of pregnancies
16. Pdiabetes:Gestation diabetes
17. UriationFreq:Frequency of urination
18. Diabetic:Yes or No

We found a total of 48 missing values in the original dataset, including 4 missing values for BMI, 42 missing values for Pregnancies, 1 missing value for Pdiabetes, and 1 missing value for Diabetic.

The boxplot in Figure 8(a) shows that the dataset contains outliers, whereas Figure 8(b) shows clean data after applying the preprocessing algorithm. In the boxplot, different features have multiple outliers data which is indicated by multiple diamond signs beside each feature and after handling the outlier, the boxplot looks like no diamond signs on each feature, which proves no outlier existed.

**Figure 8.**
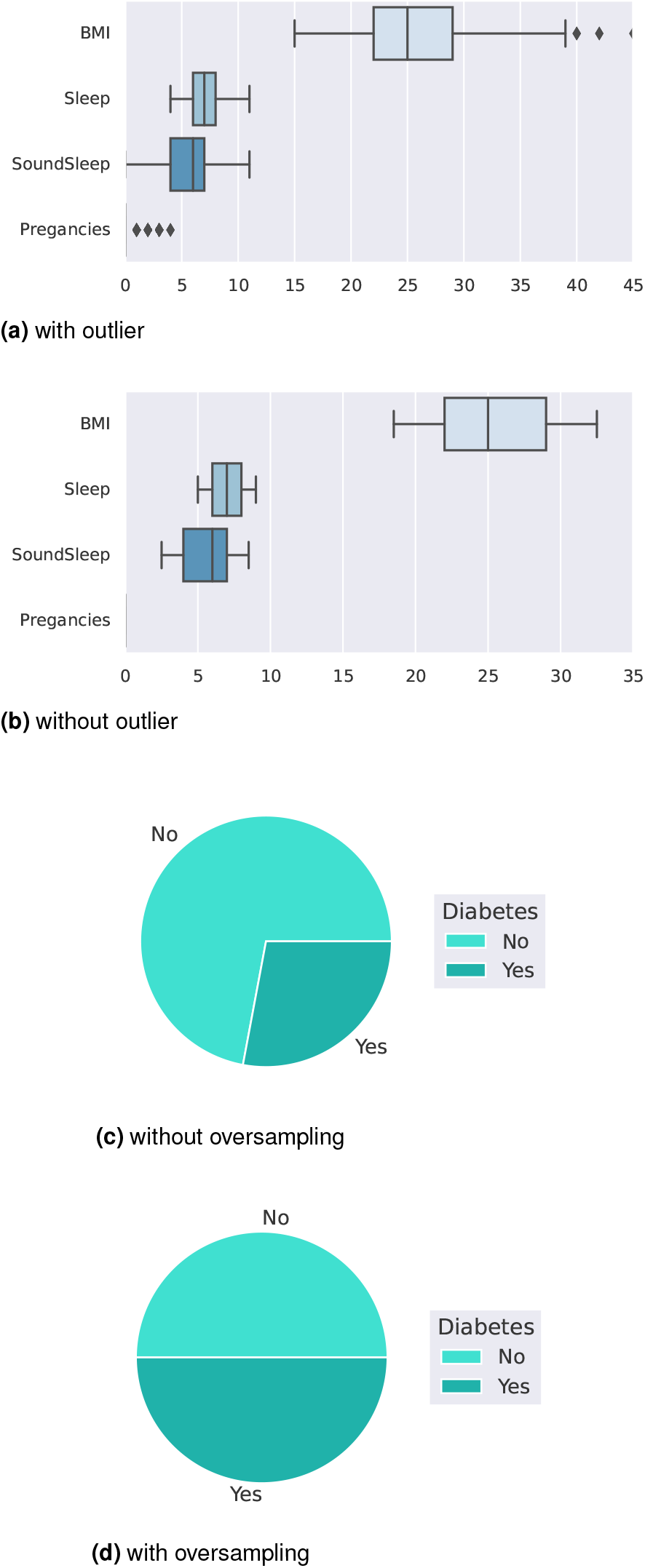
Before and after outlier removal and oversampling results

Figure 8(d) depicts the balanced dataset distribution of the original imbalance dataset 8(c). where the label ‘No’ represents no diabetes and ‘Yes’ represents diabetes. In the pie chart, we can see that it contains more portion of ‘No’ than ‘Yes’, and after random oversampling, we can see that we have an equal portion of the label ‘No’ and ‘Yes’ which ensures data is balanced now.

Figures 9(a, b) represents the accuracy and MSE of our experiments for dataset-3. The accuracy comparison before and after applying the proposal. The accuracy results of DT, RF, LR, SVM, KNN, AdB, GB and XGB are 98.54%, 97.81%, 91.11%, 91.97%, 88.32%, 91.58%, 92.63% and 99.27%, respectively. We found that, depending on the ML algorithms, accuracy performance increases from 2.25% to 10.75%. On the other hand, the MSE values of the algorithm reduced significantly, 2.98%, 2.25%, 8.63%, 10.75%, 7.66%, 3.26%, 4.31% and 3.71% for DT, RF, LR, SVM, KNN, AdB, GB and XGB respectively.

**Figure 9.**
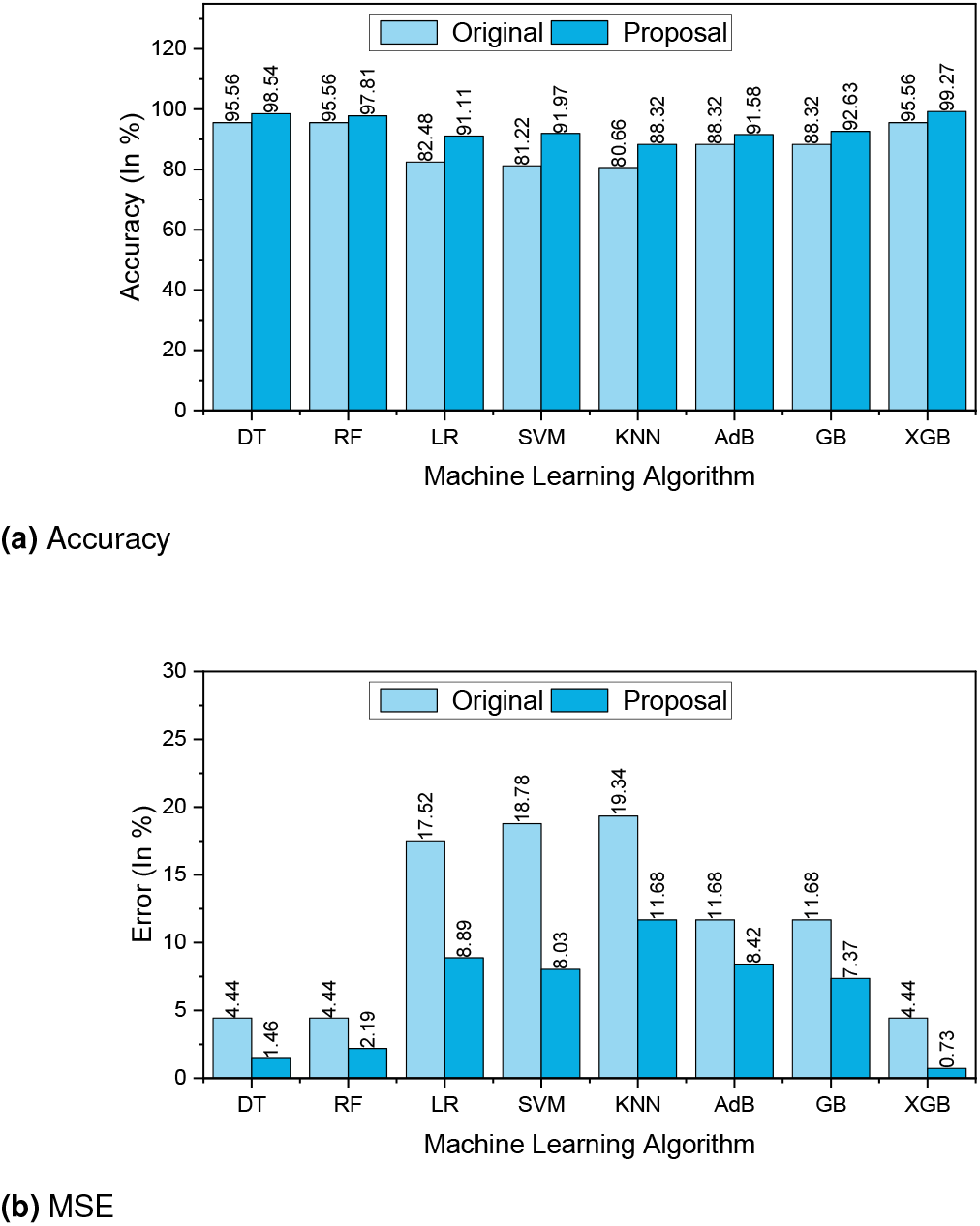
The performance Results of Dataset-3

**Figure 10.**
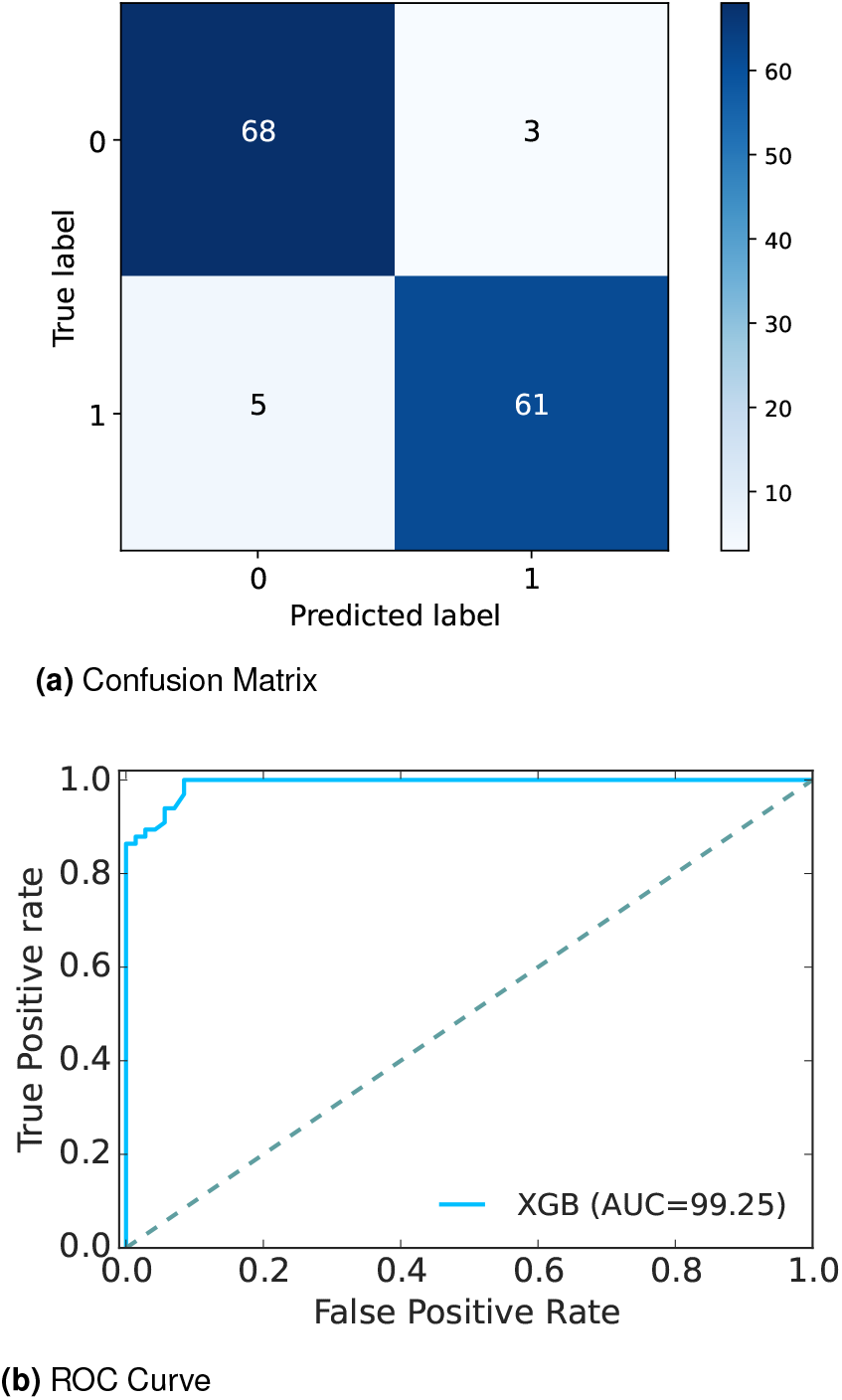
Confusion Matrix and ROC Curve of Dataset-3

The other performance metrics are summarized in Table 5. We found that, for any ML approach, the proposal can improve the precision values from 1.77% to 9.20%. An efficient data preprocessing and data balancing can improve the data quality, hence ML algorithms can accurately classify the test data. We found similar results for recall, the values increase from 2.06% to 25.34%. The F1-score also improved as expected from 1.99% to 22.74%. On the other hand, the table also indicates that due to the high performance of the proposal, the values of MAE and RMSE are reduced significantly. It is observed that MAE values reduced from 2.25% for RF to 10.75% for SVM. Similarly, RMSE reduced greatly from 5.15% to 15%.

**Table 5.**
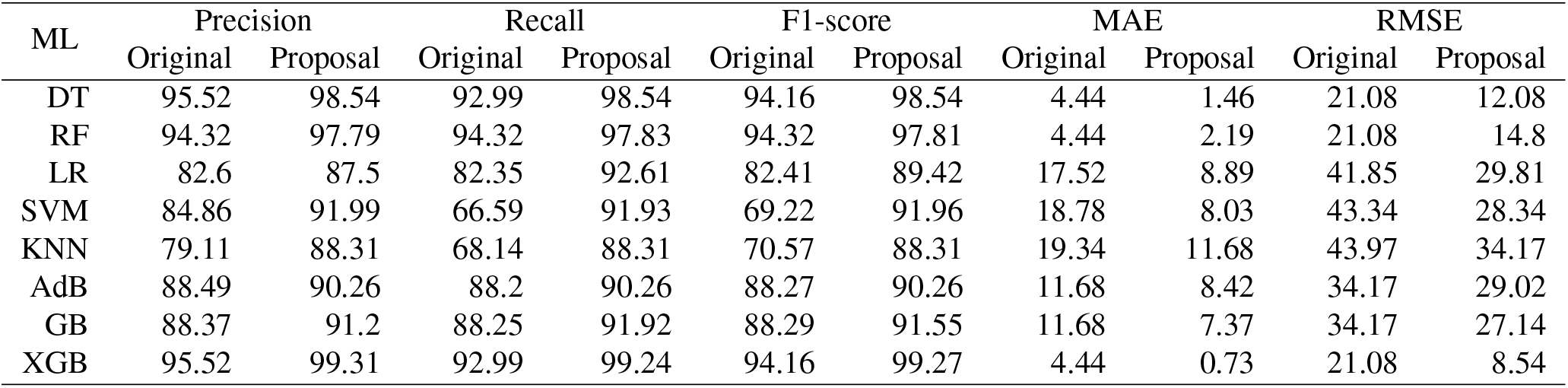
Performance result of dataset-03 (Tigga dataset)

| ML | Precision |  | Recall |  | F1-score |  | MAE |  | RMSE |  |
| --- | --- | --- | --- | --- | --- | --- | --- | --- | --- | --- |
|  | Original | Proposal | Original | Proposal | Original | Proposal | Original | Proposal | Original | Proposal |
| DT | 95.52 | 98.54 | 92.99 | 98.54 | 94.16 | 98.54 | 4.44 | 1.46 | 21.08 | 12.08 |
| RF | 94.32 | 97.79 | 94.32 | 97.83 | 94.32 | 97.81 | 4.44 | 2.19 | 21.08 | 14.8 |
| LR | 82.6 | 87.5 | 82.35 | 92.61 | 82.41 | 89.42 | 17.52 | 8.89 | 41.85 | 29.81 |
| SVM | 84.86 | 91.99 | 66.59 | 91.93 | 69.22 | 91.96 | 18.78 | 8.03 | 43.34 | 28.34 |
| KNN | 79.11 | 88.31 | 68.14 | 88.31 | 70.57 | 88.31 | 19.34 | 11.68 | 43.97 | 34.17 |
| AdB | 88.49 | 90.26 | 88.2 | 90.26 | 88.27 | 90.26 | 11.68 | 8.42 | 34.17 | 29.02 |
| GB | 88.37 | 91.2 | 88.25 | 91.92 | 88.29 | 91.55 | 11.68 | 7.37 | 34.17 | 27.14 |
| XGB | 95.52 | 99.31 | 92.99 | 99.24 | 94.16 | 99.27 | 4.44 | 0.73 | 21.08 | 8.54 |

The extensive analysis of the experiments shows that XGB outperforms others as it is a gradient-boosted decision tree solution that uses L1 and L2 regularization which leads to getting a better accuracy rate than others. So, we consider this model as our proposed model.

Graph 10(a) depicts the confusion matrix and 10(b) shows the ROC Curve for our proposed model. In the confusion matrix, the large number of TP and TN than FP and FN is a very crucial point to be a better prediction model for ML. In graph 10(a), we can see that the TP and FN rate are high and the FN and FP rates are very low which gives a better sign of diabetes prediction. The TP, TN, FP and FN rates are 49.64%, 44.53%, 3.65%, and 2.19% respectively. On the other hand, in ROC Curve the value of AUC which is near 1 is the best model for ML. In graph 10(b), we can see that the AUC value is 0.99 (99.36%) which means the positive and negative labels are almost segregated, and the model is effective.

After analyzing multiple performance indicators, we can find that among all the ML algorithms XGB outperforms others with an accuracy rate of 99.27%, 0.73% MSE rate, and 3.65% FP as well as 2.19% FN rates.

### Results of Dataset -4

Finally, the information was collected from Iraqi society, as well as the Medical City Hospital’s laboratory and Specializes Center for Endocrinology and Diabetes-AlKindy Teaching Hospital in dataset 4 (*40*). The data consist of medical information as well as laboratory analysis including age, gender, Creatinine ratio(Cr), BMI, Urea, Cholesterol, LDL, VLDL, Triglycerides and HDL Cholesterol, HBA1C, etc. The attributes of the Iraqi Medical City dataset are listed below:

1. Age:Age of the patient
2. Gender:Gender of the participant
3. Sugar Level Blood:Sugar Level of the patient
4. Cr:Creatinine ratio
5. BMI:Body Mass Index (BMI)
6. Urea:blood urea level
7. Chol:Cholesterol (Chol)
8. TG:Triglycerides (TG) level
9. HDL:HDL Cholesterol level
10. LDL:Low density lipoprotein (LDL) level
11. VLDL:Very low-density lipoprotein (VLDL) level
12. HBA1C:Average blood glucose (sugar)-Haemoglobin A1c
13. Class:Diabetic, Non-Diabetic, or Pre-Diabetic

The boxplot in Figure 11(a) shows that the dataset contains outliers, whereas Figure 11(b) shows clean data after applying the preprocessing algorithm. In the boxplot, different features have multiple outliers data which is indicated by multiple diamond signs beside each feature and after handling the outlier the boxplot looks like no diamond signs on each feature which proves no outlier existed.

**Figure 11.**
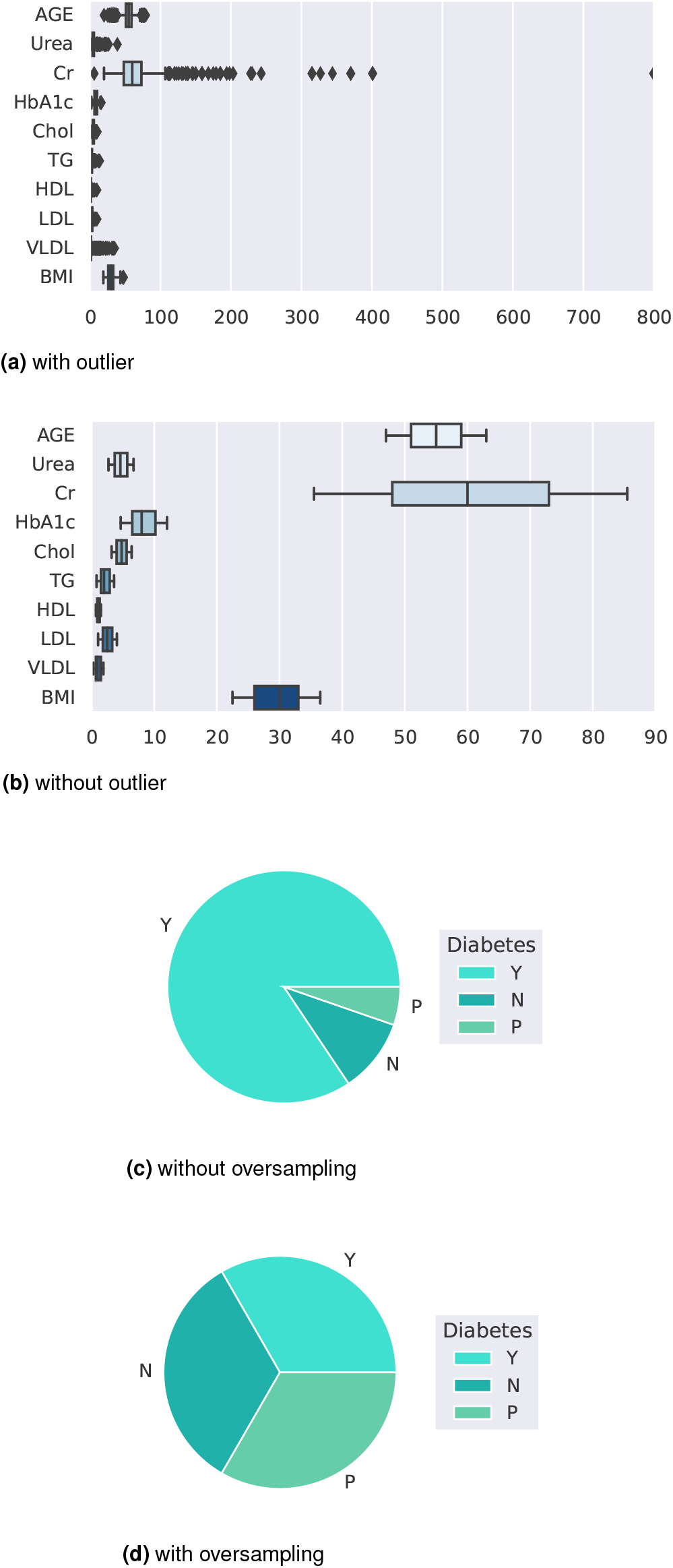
Before and after outlier removal and oversampling results

Figure 11(d) depicts the balanced dataset distribution of the original imbalance dataset 11(c). where the label ‘Y’ represents diabetic, ‘N’ represents non-diabetic and ‘P’ represents pre-diabetic. In the pie chart, we can see that it contains more portion of ‘Y’ than ‘P’ and ‘N’, and after random oversampling, we can see that we have an equal portion of all the labels which ensures data is balanced now.

The research specifically aims to develop an ML model for classifying diabetes, focusing on the task of assigning diabetes labels (diabetes or no diabetes) using various diabetes datasets. While it may seem odd to include HbA1c as an input variable in this dataset (dataset-4), we considered it relevant for our classification task. The inclusion of HbA1c as a feature helps the model learn patterns and relationships between other variables and the presence of diabetes. By including this feature, we aim to capture additional information that may contribute to the accurate classification of diabetes. We understand that the close-to-perfect performance of the model might raise suspicions. However, we assure you that our research was conducted rigorously, following standard practices and using appropriate evaluation metrics.

Figures 12(a, b) represent the accuracy and MSE of our experiments for dataset-4. The accuracy comparison before and after applying the proposal. The accuracy results of DT, RF, LR, SVM, KNN, AdB, GB and XGB are 100%, 99.60%, 95.65%, 96.84%, 95.65%, 99.21%, 99.60% and 99%, respectively. We found that, depending on the ML algorithms, accuracy performance increases from 0.19% to 9.84%. On the other hand, the MSE values of the algorithm reduced significantly, 2%, 1.6%, 11.09%, 21.84%, 18.28%, 3.71%, 7.6% and 0.19% for DT, RF, LR, SVM, KNN, AdB, GB and XGB respectively.

**Figure 12.**
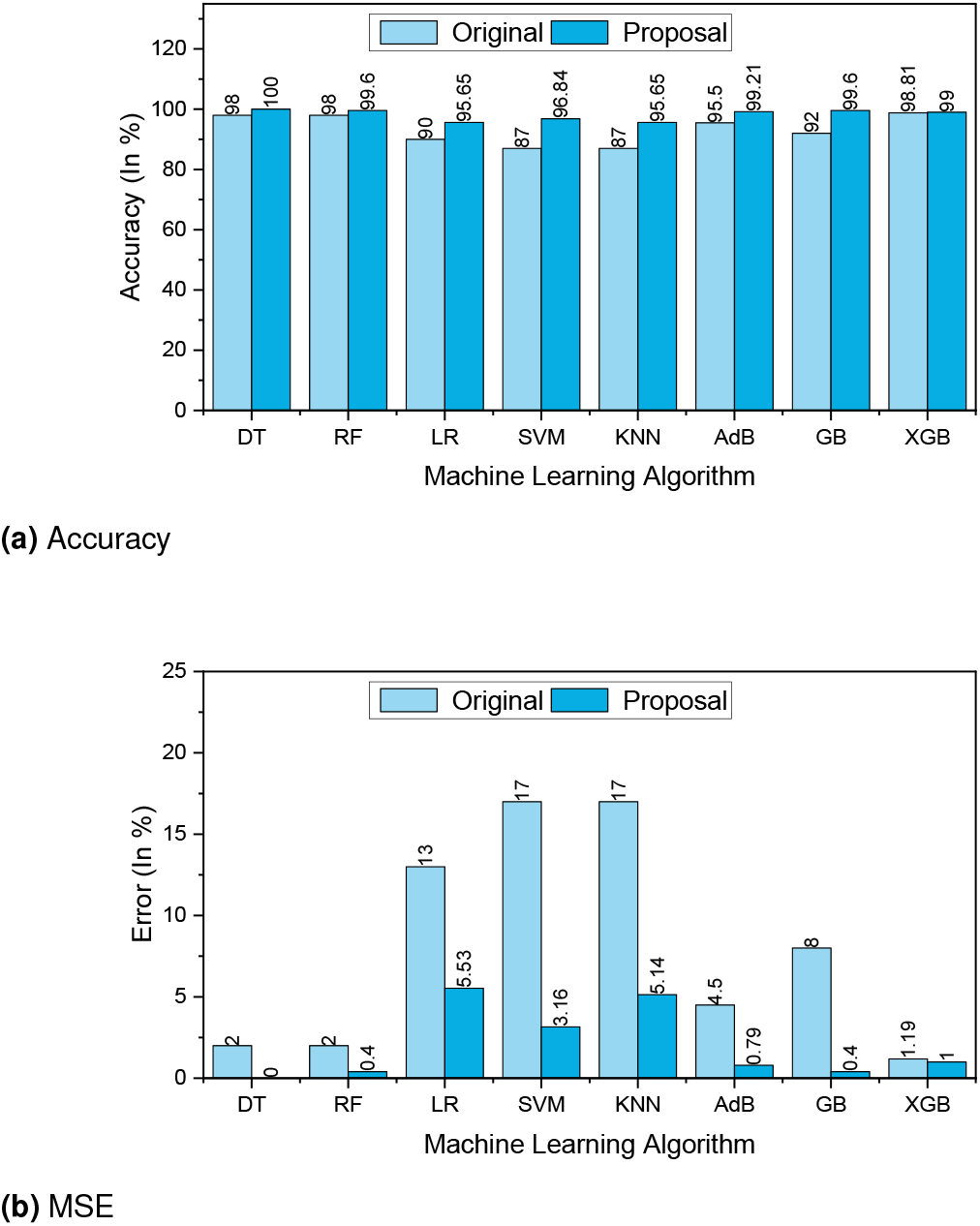
The performance Results of Dataset-4

**Figure 13.**
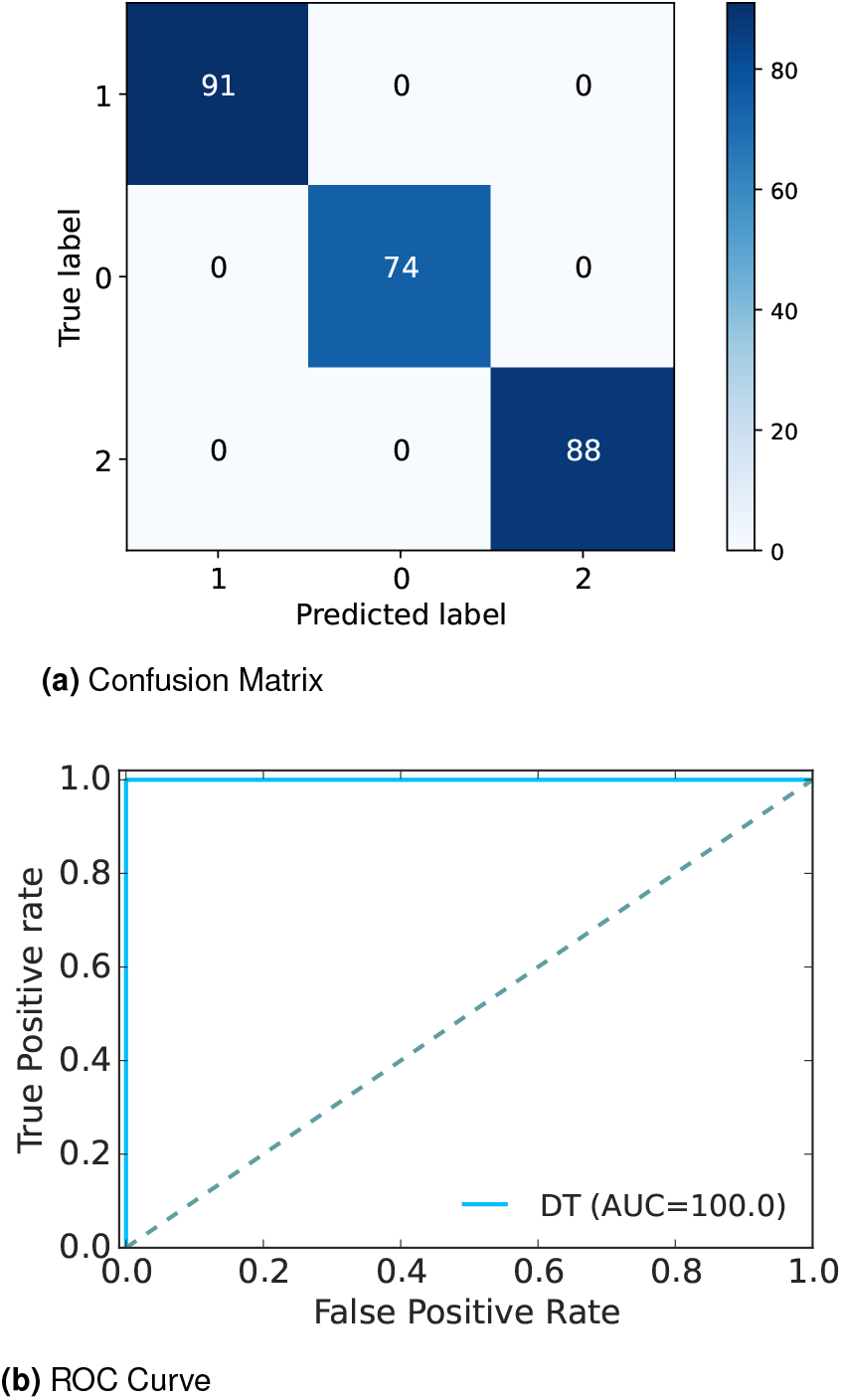
Confusion Matrix and ROC Curve for RF of Dataset-4

**Figure 14.**
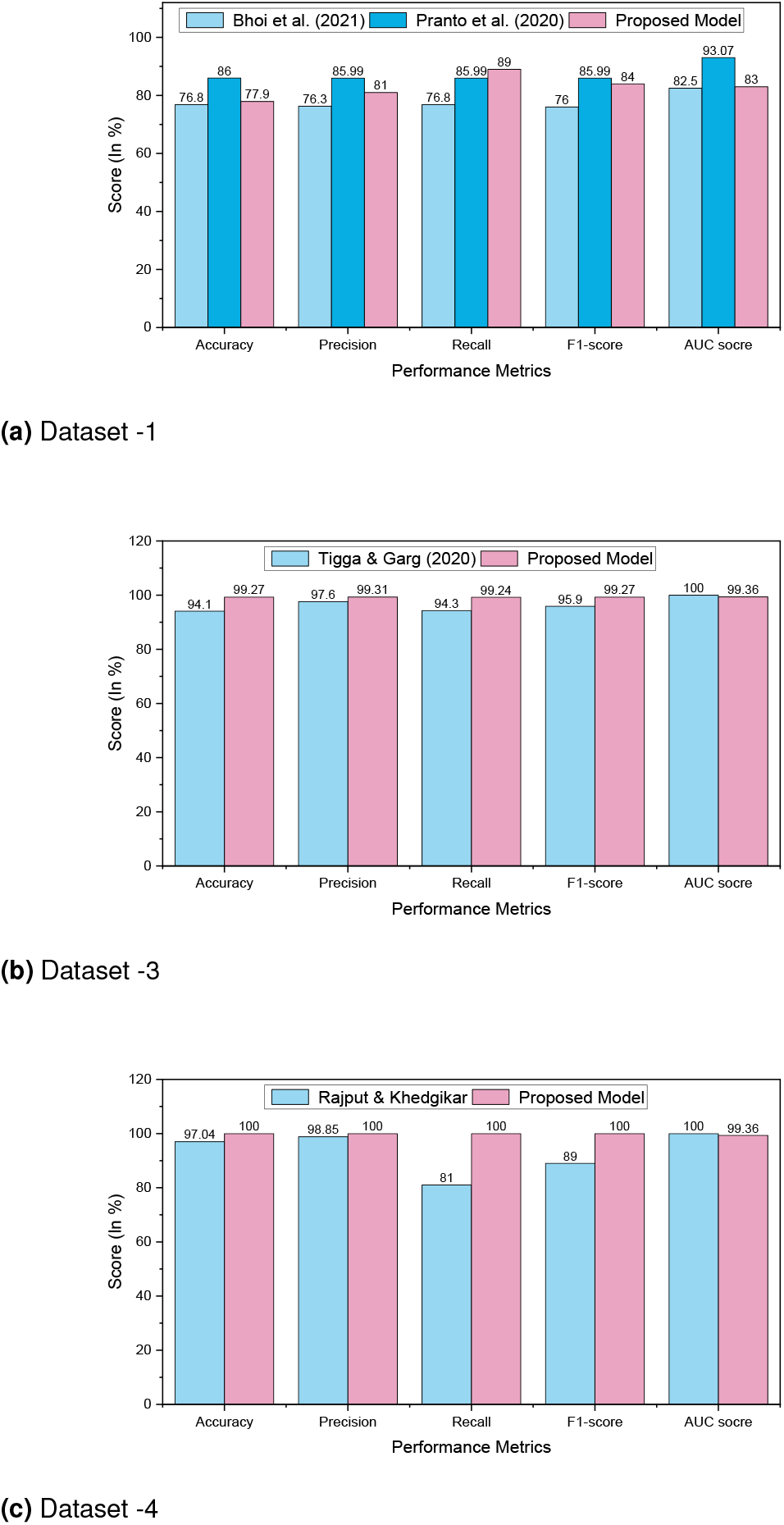
Comparison analysis of diabetes prediction for dataset -1, dataset -3, and dataset -4

The other performance metrics are summarized in Table 6. We found that, for any ML approach, the proposal can improve the precision values of 0.67% to 46.88%. An efficient data preprocessing and data balancing can improve the data quality, hence ML algorithms can accurately classify the test data. We found similar results for recall, the values increased from 0.83% to 47.59%. The F1 score also improved as expected from 0.42% to 46.52%. On the other hand, the table also indicates that due to the high performance of the proposal, the values of MAE and RMSE are reduced significantly. It is observed that MAE values reduced from 0.19% for XGB to 13.84% for SVM. Similarly, RMSE reduced greatly from 0.89% to 32.22%. XGB 98.94 99.61 97.22 98.65 98.35 98.77 1.19 1 10.89 10

**Table 6.** Performance result of dataset-04 (Mendeley dataset)

| ML | Precision |  | Recall |  | F1-score |  | MAE |  | RMSE |  |
| --- | --- | --- | --- | --- | --- | --- | --- | --- | --- | --- |
|  | Original | Proposal | Original | Proposal | Original | Proposal | Original | Proposal | Original | Proposal |
| DT | 93.19 | 100 | 96.44 | 100 | 94.61 | 100 | 2 | 0 | 14.14 | 0 |
| RF | 92.94 | 99.63 | 96.83 | 99.55 | 94.45 | 99.58 | 2 | 0.4 | 14.14 | 6.29 |
| LR | 67.48 | 96.17 | 70.63 | 95.3 | 68.46 | 95.55 | 13 | 5.53 | 43.59 | 28.12 |
| SVM | 53.26 | 97.16 | 48.81 | 96.4 | 50.63 | 96.65 | 17 | 3.16 | 50 | 17.78 |
| KNN | 49.22 | 96.1 | 48.81 | 95.21 | 48.97 | 95.49 | 17 | 5.14 | 50 | 25.92 |
| AdB | 79.44 | 99.26 | 98.27 | 99.1 | 85 | 99.17 | 4.5 | 0.79 | 21.21 | 8.89 |
| GB | 86.88 | 99.64 | 89.68 | 99.55 | 88.14 | 99.59 | 8 | 0.4 | 28.28 | 6.29 |
| XGB | 98.94 | 99.61 | 97.22 | 98.65 | 98.35 | 98.77 | 1.19 | 1 | 10.89 | 10 |

The extensive analysis of the experiments shows that DT outperforms others since the ability to capture relevant decision-making information from the available dataset is the most important feature of the decision tree which leads to higher accuracy. So, we consider this model as our proposed model.

Graph 13(a) depicts the confusion matrix and 13(b) shows the ROC Curve for our proposed model. In the confusion matrix, the large number of TP and TN than FP and FN is a very crucial point to be a better prediction model for ML.

In graph 13(a), we can see that the TP and FN rates are high and FN and FP rates are very low which gives a better sign of diabetes prediction. The TP, TN, FP and FN rates are 35.97%, 64.03%, 0.0%, 0.0%; 29.25%, 70.75%, 0.0%, 0.0%; 34.78%, 65.22%, 0.0%, 0.0% for N, Y,and P class respectively. On the other hand, in the ROC, the value of AUC which is near 1 is the best model for ML. In the graph 13(b), we can see that the AUC value is 1 (100%) which means the positive and negative labels are completely segregated, and the model is as effective as it can be.

After analyzing multiple performance indicators, we can find that among all the ML algorithms DT outperforms others with an accuracy rate of 100%, 0% MSE rate, and 0% FP as well as FN rates.

## Discussion

In this study, we conducted a comprehensive analysis of ML models for diabetes detection using four distinct datasets: Pima Indian, Austin Public, Tigga dataset, and Mendeley. The performance metrics were evaluated to assess the effectiveness of the models in accurately identifying individuals at risk of diabetes as shown in Table 7.

**Table 7.**
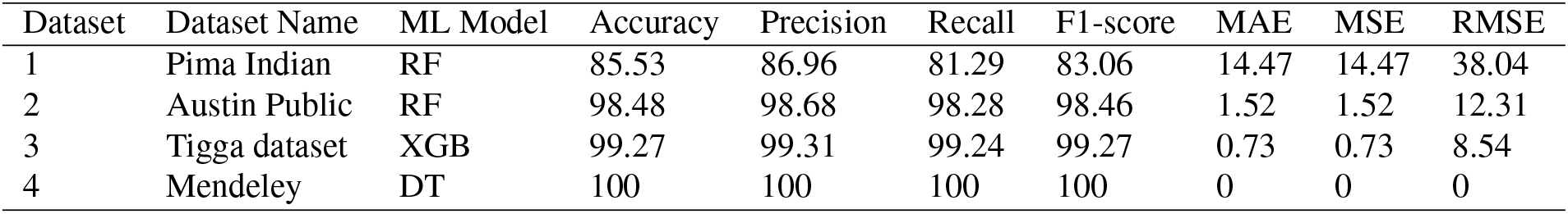
Best performance analysis for each dataset.

| Dataset | Dataset Name | ML Model | Accuracy | Precision | Recall | F1-score | MAE | MSE | RMSE |
| --- | --- | --- | --- | --- | --- | --- | --- | --- | --- |
| 1 | Pima Indian | RF | 85.53 | 86.96 | 81.29 | 83.06 | 14.47 | 14.47 | 38.04 |
| 2 | Austin Public | RF | 98.48 | 98.68 | 98.28 | 98.46 | 1.52 | 1.52 | 12.31 |
| 3 | Tigga dataset | XGB | 99.27 | 99.31 | 99.24 | 99.27 | 0.73 | 0.73 | 8.54 |
| 4 | Mendeley | DT | 100 | 100 | 100 | 100 | 0 | 0 | 0 |

On the Pima Indian dataset, the Random Forest (RF) model achieved an accuracy of 85.53%. The precision and recall were 86.96% and 81.29%, respectively, resulting in an F1-score of 83.06%. The Mean Absolute Error (MAE) and Mean Squared Error (MSE) were 14.47, and the Root Mean Squared Error (RMSE) was 38.04. For the Austin Public dataset, the RF model exhibited exceptional performance with an accuracy of 98.48%. The precision, recall, and F1-score values were 98.68%, 98.28%, and 98.46%, respectively. The model demonstrated low errors with an MAE of 1.52, MSE of 1.52, and RMSE of 12.31. The Tigga dataset showed outstanding results with the XGBoost (XGB) model achieving an accuracy of 99.27%. The precision, recall, and F1-score values were 99.31%, 99.24%, and 99.27%, respectively. The model’s errors were minimal with an MAE of 0.73, MSE of 0.73, and RMSE of 8.54. Remarkably, the Decision Tree (DT) model achieved perfect performance on the Mendeley dataset, attaining an accuracy, precision, recall, and F1-score of 100%. Additionally, the model exhibited zero errors with an MAE, MSE, and RMSE of 0.

Interestingly, the DT model achieves perfect performance on the Mendeley dataset, with accuracy, precision, recall, and F1-score all reaching 100%. This remarkable result suggests that the features within the Mendeley dataset may be well-suited for decision tree-based classification, possibly due to the dataset’s inherent structure or the nature of the variables involved. It is worth noting that while the DT model demonstrates flawless performance on this particular dataset, its generalization to other datasets may vary, warranting further investigation into its robustness across different data domains. Moreover, the low error metrics (MAE, MSE, RMSE) observed across all models and datasets indicate the models’ capability to make accurate predictions with minimal deviation from the actual values. These findings emphasize the reliability of ML algorithms in diabetes detection tasks and underscore their potential utility in clinical settings for early risk assessment and intervention.

Furthermore, the results of this study highlight the efficacy of ML models in diabetes detection across diverse datasets. While certain models excel in specific contexts, the overall performance underscores the promise of ML techniques in augmenting traditional diagnostic approaches and improving patient outcomes in diabetes management. Further research is warranted to explore the generalizability of these models across larger and more diverse populations, as well as their integration into clinical practice for personalized healthcare delivery.

The results indicate that the ML models can effectively detect diabetes. The RF model showed good performance on the Pima Indian dataset, while both the RF model on the Austin Public dataset and the XGB model on the Tigga dataset demonstrated excellent performance. The DT model exhibited perfect performance on the Mendeley dataset. These findings highlight the potential of ML in accurate diabetes detection, providing a valuable tool for early intervention and improved patient outcomes.

In our research, we included multiple datasets in our analysis to provide a comprehensive evaluation of the performance of ML models for diabetes detection. Each dataset represents a distinct population or data source, allowing us to assess the generalizability of the models across diverse scenarios. We evaluated the models on multiple datasets to gain insights into their strengths and limitations in different contexts. This approach helps experimenters understand the robustness of the models and identify potential challenges or biases that may arise when applying them to real-world scenarios. Additionally, it enables researchers to make informed decisions about which models are most suitable for specific datasets or patient populations. Regarding the conclusions, we acknowledge that the original discussion did not sufficiently elaborate on the insights gained from the extensive set of benchmarks. In light of the reviewer’s comment, we will revise the conclusion section to provide a more comprehensive analysis of the results and their implications. We will discuss the key findings from each dataset, highlight the factors that contributed to successful performance, and address the challenges and considerations experimenters should be aware of when deploying these models in practice. We will also emphasize any new information or novel observations that emerged from our study. Although some previous research has examined ML models for diabetes detection, our study contributes by analyzing a diverse range of datasets and comparing the performance of multiple models. This allows us to provide a more comprehensive understanding of the strengths and weaknesses of different algorithms and their applicability in various scenarios.

Furthermore, a comparison analysis is illustrated in Table 8 and in Bar chart 14, where we can see that our proposed approach outperforms others, which proves the better prediction models. In dataset -1, RF outperforms other ML models with an accuracy rate of 86%. Similarly, for dataset -2, dataset -3, and dataset -4, RF, XGB, and DT outperform other ML models with an accuracy rate of 98.48%, 99.27%, and 100% respectively. The higher accuracy rate of diabetes predictions proves the robustness of our proposed model.

**Table 8.**
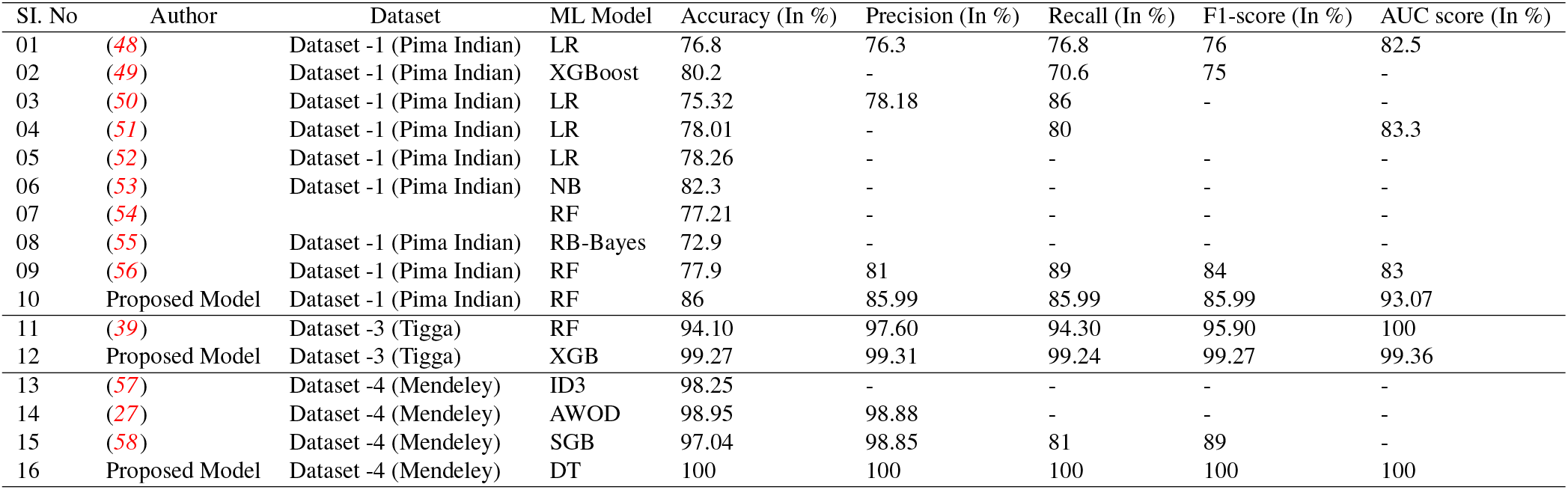
Comparison analysis of diabetes prediction for dataset -1, dataset -3, and dataset -4.

The ML models RF, XGB, and DT exceed the performance indicators of other algorithms as well as research for dataset -1, dataset -2, dataset -3, and dataset -4. The proper efficient preprocessing upgrades the quality of data that helps to enhance the outcomes for various datasets. Yet, the RF, XGB, and DT algorithms had enhanced accuracy, and it is encouraged that they can be employed in the clinical categorization and prognosis of diabetes for greater performance.

### Novelty and Significance

This study acknowledges the well-established nature of optimizing preprocessing and oversampling techniques within the specific context of diabetes prediction using ML models. Our contribution lies in carefully implementing and evaluating these methods within the diabetes detection framework.

#### Context-specific application

Although oversampling techniques are not new, their effectiveness and impact on diabetes prediction may vary depending on the dataset and the specific problem being addressed. By applying these techniques in the context of diabetes detection, we aimed to explore their potential benefits and address the challenges associated with imbalanced class problems specific to this domain.

#### Comparative analysis

Our study not only employed oversampling techniques but also performed a comparative analysis of different approaches for handling imbalanced class problems, such as undersampling and synthetic data generation. This analysis aimed to identify the most suitable technique for diabetes prediction and provide insights into the trade-offs and limitations of each approach.

#### Rigorous evaluation

We implemented a robust evaluation methodology by employing *k*-fold cross-validation. This approach ensures that our models are thoroughly tested on different subsets of the dataset, reducing the risk of overfitting and providing a more reliable assessment of their performance.

While our study introduces the experimental evaluation techniques, we believe that the methodological and clinical insights gained from our rigorous analysis contribute to the field of diabetes prediction. By carefully implementing and evaluating oversampling techniques within the diabetes detection framework, we aim to provide practical guidance for researchers and practitioners working on similar problems.

The significance of this study lies in its contributions to the field of diabetes research and its potential impact on clinical practice.

#### 1. Improved diagnostic accuracy

By developing an optimized data preprocessing pipeline and implementing advanced ML techniques, our study enhances the accuracy of diabetes prediction. This can assist healthcare professionals in making more precise and timely diagnoses, leading to better patient outcomes.

#### 2. Personalized diabetes management

The robust models developed in this study have the potential to enable personalized diabetes management. By accurately predicting diabetes prognosis, healthcare providers can tailor treatment plans and interventions to individual patients, optimizing their care and reducing the risk of complications.

#### 3. Early intervention and prevention

Early detection of diabetes is crucial for effective intervention and prevention. Our study contributes to the development of predictive models that can identify individuals at high risk of developing diabetes. This enables early intervention strategies, such as lifestyle modifications and targeted preventive measures, to be implemented, reducing the burden of the disease.

#### 4. Advancement in ML techniques

Through the exploration and evaluation of various ML algorithms and preprocessing techniques, our study contributes to the advancement of the field. The insights gained from this research can inform the development of more robust and interpretable machine-learning models for diabetes prediction and prognosis.

Overall, this study holds significant implications for clinical practice, offering improved diagnostic accuracy, personalized management strategies, early intervention opportunities, and advancements in ML techniques. The findings have the potential to enhance diabetes care, contribute to preventive healthcare, and ultimately improve patient outcomes.

## Conclusion

In this research, we have conducted a comprehensive study on diabetes detection using machine learning techniques, aiming to underscore both the scientific value added by our work and the applicability of our findings in clinical practice. Our contributions encompass the outcome of an optimized preprocessing pipeline, addressing dataset imbalance, preventing overfitting, and demonstrating superior performance through extensive experimentation. We have rigorously evaluated various machine learning models on four different datasets: Pima Indian, Austin Public, Tigga, and Mendeley. Our results showcase notable improvements in accuracy, precision, recall, and F1-score metrics compared to existing methods.

Specifically, the RF model achieved an accuracy of 85.53% on the Pima Indian dataset, with balanced precision and recall values. On the Austin Public dataset, the Random Forest model excelled with an exceptional accuracy of 98.48%, along with high precision, recall, and F1-score values. The XGBoost model demonstrated outstanding performance on the Tigga dataset, achieving an accuracy of 99.27% with minimal errors in predictions. Notably, the DT model achieved perfect accuracy and precision on the Mendeley dataset, indicating flawless classification of diabetes instances. Our study reveals significant improvements over existing methods, with accuracy rates ranging from 86% to 100% across different datasets. Specifically, our suggested method outperforms previous works by 4.95% to 12.15% for dataset-1, 0% to 8.74% for dataset-2, 2.25% to 10.75% for dataset-3, and 0.19% to 9.84% for dataset-4.

However, it’s essential to acknowledge the limitations of our study. Further investigation is needed to assess the generalizability of our approach to diverse datasets, feature selection, ensemble models and DL techniques. Additionally, the lack of interpretability in ML models poses a challenge in understanding the underlying factors driving predictions.

In conclusion, our study highlights the need for further research to address limitations and enhance the reliability and applicability of the proposed approach for diabetes detection using ML. Moving forward, potential avenues for future research include:

### 1. Feature Selection

Exploring advanced feature selection techniques to improve the efficiency and accuracy of diabetes detection models.

### 2. Ensemble Models

Investigating the integration of ensemble learning techniques to combine multiple models for enhanced predictive performance.

### 3. Deep Learning Algorithms

Exploring the application of deep learning algorithms, such as convolutional neural networks (CNNs) and recurrent neural networks (RNNs), to improve the prediction accuracy of diabetes detection models.

By pursuing these future directions, we aim to advance the field of diabetes detection using ML and contribute to improved healthcare outcomes.

## Declarations

### Guarantor

Md. Alamin Talukder.

### Ethics approval

Not applicable

### Consent to participate

Not applicable

### Consent to Publish

Not applicable

### Availability of data and materials

The UCI ML Diabetes dataset-1 is available on https://archive.ics.uci.edu/ml/datasets/diabetes; The Austin Public Health Diabetes dataset-2 is available on https://data.austintexas.gov/Health-and-Community-Services/Austin-Public-Health-Diabetes-Self-Manageme-AEldbuercti/48iy-4sbg; The Type 2 Diabetes dataset-3 is available on https://www.kaggle.com/datasets/tigganeha4/diabetes-dataset-2019; The Mendeley Diabetes dataset-4 is available on https://data.mendeley.com/datasets/wj9rwkp9c2/1

## Authors’ contributions

Md. Alamin Talukder: Conceptualization, Data Curation, Methodology, Software, Resource, Visualization, Formal Analysis, Writing–original draft and Review & Editing; Md. Manowarul Islam, Md Ashraf Uddin, Arnisha Akther and Mohammad Ali Moni: Methodology, Visualization, Validation, Investigation, Writing–Review & Editing; Majdi Khalid, Mohsin Kazi: Supervision, Formal Analysis, Visualization, Investigation, Writing–Review & Editing.

